# Disruption of bacteriophage integration site promotes rapid diversification of multicellular traits in *Bacillus subtilis*

**DOI:** 10.1101/2025.08.01.668206

**Authors:** Maja Popović, Tomaž Accetto, Yasmine Dergham, Romain Briandet, Iztok Dogša, Anna Dragoš

## Abstract

Certain bacteria are known for their remarkable genetic and phenotypic diversity, as well as rapid morphological diversification during evolution experiments. An example is *Bacillus subtilis*, which can switch motility, biofilm or antagonistic interaction patterns either as a result of spontaneous mutations or due to changes in prophage elements. Prophages can integrate into conserved and functional loci, disrupting host genes and modulating their phenotypic traits. In *B. subtilis*, the SPβ prophage integrates into the sporulation-associated gene *spsM*, whose reversible inactivation during lysogeny has also been implicated in modulating biofilm formation and host development. Here, we investigated the evolutionary and phenotypic consequences of *spsM* disruption through both SPβ integration and artificial mutagenesis (*spsM::kan*) in *B. subtilis* natural isolates.

We observed that *spsM::kan* mutants frequently developed spontaneous mutations, particularly in *swrA* and *comP*, key regulators of swarming motility and biofilm development. These mutations reproducibly gave rise to altered colony morphotypes and impaired surface motility, suggesting strong selection for loss-of-function mutations in regulatory genes under laboratory conditions. In contrast, SPβ lysogens exhibited minimal mutational diversification, indicating that dynamic prophage excision may preserve genomic stability. In this work *spsM* inactivation alone did not significantly impair biofilm formation or motility. However, we reveal a novel role of prophage integration sites as possible evolutionary hotspots that influence genome integrity and adaptive potential. This work highlights the interplay between prophage integration, host genome architecture, and the selective pressures shaping bacterial multicellular communities.

**IMPORTANCE:** Prophages, defined as viruses integrated into bacterial genomes, can reshape bacterial physiology and evolution. Previous studies suggested that disruption of an integration site (*spsM*) by the SPβ prophage impairs biofilm formation in *Bacillus subtilis*, yet the functional basis for this remained unclear. Here, we show that *spsM* disruption across diverse natural isolates promotes the rapid emergence of spontaneous mutations in key regulatory genes like *swrA* and *comP*, which do influence biofilm morphology and motility. Strikingly, when *spsM* is disrupted by a prophage capable of precise excision, such diversification is minimized, indicating a protective role for dynamic prophage integration. These findings reconcile data from earlier work and identify prophage integration sites as evolutionary hotspots, affecting host genome stability. This has broader implications for how we understand the genetic basis of microbial adaptation and the evolutionary roles of prophages.

## INTRODUCTION

Bacteria adapt rapidly to new ecological conditions, a phenomenon that makes them an ideal experimental model for tracking evolution in real time and decoding molecular evolutionary patterns (Kovács & Dragoš, 2019; McDonald, 2019). In structured environments such as solid media, multiple evolving lineages can coexist for extended periods due to spatial structure and niche separation (Kuo et al., 2025; M. Martin et al., 2016). In some cases, bacterial diversification becomes macroscopically visible, as outgrowths emerging from original colony outpace the ancestor in spreading across solid media (Baym et al., 2016; Kim et al., 2016).

Certain bacterial species exhibit exceptionally high levels of genetic diversity (Dewar et al., 2024), which is often reflected in striking phenotypic variation (Stefanic, 2020). A well-studied example is *Bacillus subtilis*, a soil-dwelling, spore-forming bacterium well known for its developmental plasticity, including its ability to form biofilms (Arnaouteli et al., 2021) and undergo sporulation (Tan & Ramamurthi, 2014). In biofilm evolution experiments, *B. subtilis* has been shown to diversify into distinct morphotypes within just a few days (Dragoš et al., 2017; Leiman et al., 2014; Richter et al., 2018). This rapid diversification is also commonly observed during laboratory domestication and even through routine strain-sharing practices, often resulting in inconsistent phenotypes among strains thought to be genetically identical (Gallegos-Monterrosa et al., 2016). Most reported mutations are those that are associated with an easily recognizable phenotype, such as changes in motility, changes in biofilm formation or emergence of antagonistic interactions (Dragoš, Priyadarshini, et al., 2021; M. Martin et al., 2017). Notably, the latter two have been associated with the acquisition of new prophage elements, which are particularly prevalent in *B. subtilis* (Stefanič et al., 2024).

Many natural isolates of *B. subtilis* carry multiple prophages (Stefanič et al., 2024) with members of the *Spbetavirus* genus, particularly the SPβ phage, being among the most thoroughly characterized (Kohm et al., 2022; Kohm & Hertel, 2021). SPβ is a temperate siphovirus phage, approximately 130 kb in size (Abe et al., 2014). It was previously shown that SPβ can rapidly alter host phenotype by inducing the production of the bacteriocin sublancin, which contributes to antagonism against ancestral prophage-free variants (Dragoš, Andersen, et al., 2021). Beyond this antagonistic effect, SPβ also modifies its host through site-specific integration into the *spsM* gene (formerly ypqP), a locus reported to be involved in sporulation and biofilm development (Abe et al., 2014; T. Dubois et al., 2020; Sanchez-Vizuete et al., 2015). The *spsM* gene encodes a polysaccharide synthase involved in the synthesis of legionaminic acid (Leg), a key component of the spore crust that enhances dispersibility by increasing spore surface hydrophilicity (T. Dubois et al., 2020).

Integration of SPβ into *spsM* results in temporary gene disruption and is reversed through site-specific excision prior to sporulation (Abe et al., 2014). This reversible integration-excision mechanism, known as active lysogeny, has also been described in other bacterial species, such as *Clostridioides difficile*, *Listeria monocytogenes*, and *Streptococcus pneumoniae*, as well as at additional chromosomal loci in *B. subtilis* (Feiner et al., 2015). Active lysogeny represents a sophisticated prophage-host relationship where the prophage excises under specific physiological conditions to restore host gene function. During sporulation, SPβ is known to excise precisely from the *spsM* locus in the mother cell, restoring gene integrity and enabling the expression of *spsM*-dependent functions during late developmental stages (Abe et al., 2014). This process mirrors similar prophage-like excisions in other loci, such as the *skin* element, which is essential for reconstituting the *sigK* gene involved in sporulation (Kunkel et al., 1990; Stragier et al., 1989).

Regulation of host genes through active lysogeny raises important questions about genome stability and the long-term consequences for both the host and the phage itself. Recent studies have suggested that SPβ integration into the *spsM* locus may impair biofilm formation, potentially enhancing phage horizontal transmission within the bacterial population (Floccari & Dragoš, 2023; Sanchez-Vizuete et al., 2015). Biofilms are multicellular bacterial aggregates, where bacteria are embedded in an extracellular matrix (ECM), essential for environmental resilience and interspecies interactions (Arnaouteli et al., 2021). They can also block external phage attacks or their transmission within the population (Bond et al., 2021; Winans et al., 2024; Zhang et al., 2020). Given the ecological relevance of biofilms and the complex social traits they entail (Kovács & Dragoš, 2019; Vlamakis et al., 2008), prophage integration into biofilm-associated loci could represent an adaptive strategy that enables phages to manipulate host development and behavior. Such insertions can modulate biofilm architecture, alter sporulation dynamics, and influence competitive interactions with closely related strains—ultimately shaping both host fitness and community structure. (Floccari & Dragoš, 2023).

To better understand the eco-evolutionary consequences of *spsM* disruption in *B. subtilis*, we examined its functional outcomes following either SPβ phage integration or targeted mutagenesis. Specifically, we aimed to test whether *spsM* inactivation by SPβ represents a prophage-driven mechanism that hinders host biofilm formation. Although our findings did not support a direct role for *spsM* in biofilm development across all tested backgrounds, they suggest that prophage integration and excision at this site could play a subtler role in maintaining genome stability. These insights reinforce the idea that prophage integration sites serve as evolutionary hotspots with far-reaching implications for both host and phage biology. Our study highlights the importance of exploring prophage-host interactions beyond classical lysis-lysogeny paradigms. As we continue to unravel the complexity of phage regulatory switches and their integration targets, understanding how these elements influence host genome stability, adaptability, and behavior will be crucial for advancing our knowledge of microbial evolution and ecology.

## MATERIALS AND METHODS

### Bacterial strains and growth conditions

Bacterial strains used and constructed in this study are presented in Table 1. Plasmids and oligonucleotides used for strain construction and Sanger sequencing are presented in Table 2 and Table 3, respectively. Accession numbers of genomes analysed with whole-genome sequencing are presented in Table 4. All the strains used were first transferred form −80°C 10% glycerol stock cultures into 5 ml liquid lysogeny broth (LB, Condalab, Spain) media and incubated at 37°C and 220 rpm for 16 hours. The cultures were then used for further experiments.

**Table 1:** *B. subtilis* and *Escherichia coli* strains used in experiments or as source of gDNA and plasmids for strain construction.

| Strain label | Background and genotype | Source or reference |
| --- | --- | --- |
| ***B. subtilis*** |  |  |
| DS215 | 3610 *swrA::tet* | (Kearns et al., 2004) |
| BKE31690 | 168 *comP::ery* | (Koo et al., 2017) |
| MB8_B7 | *spsM::SPβ* (natural lysogen) | (Kiesewalter et al., 2020) |
| DTUB222 | *amyE::mkate (spec)* | (Dragoš, Andersen, et al., 2021) |
| DTUB43 | *amyE::gfp (cm)* | (Dragoš, Andersen, et al., 2021) |
| PS-216 wt | wild type | (Stefanic & Mandic-Mulec, 2009) |
| PS-216 *spsM::kan* | *spsM::kan* | This work |
| ***B. subtilis* P9_B1** |  |  |
| wt | wild type | (Kiesewalter et al., 2020) |
| *spsM::kan* | *spsM::kan* | (Dragoš, Andersen, et al., 2021) |
| *spsM::kan amyE::spsM* | *spsM::kan amyE::spsM (spec)* | This work |
| Lys SPβ | *spsM::SPβ* | (Dragoš, Andersen, et al., 2021) |
| fluffy | *spsM::kan swrA* | This work |
| *swrA::tet* | *swrA::tet* | This work |
| *spsM::kan swrA::tet* | *spsM::kan swrA::tet* | This work |
| fluffy GFP | *spsM::kan swrA amyE::gfp (cm)* | This work |
| fluffy mKate | *spsM::kan swrA amyE::mkate (spec)* | This work |
| *swrA::tet* GFP | *swrA::tet amyE::gfp (cm)* | This work |
| *swrA::tet* mKate | *swrA::tet amyE::mkate (spec)* | This work |
| *spsM::kan swrA::tet* GFP | *spsM::kan swrA::tet amyE::gfp (cm)* | This work |
| *spsM::kan swrA::tet* mKate | *spsM::kan swrA::tet amyE::mkate (spec)* | This work |
| *spsM::kan amyE::mKate* | *spsM::kan amyE::mkate (spec)* | This work |
| ***B. subtilis* NDmed** |  |  |
| wt | wild type | (D. J. H. Martin et al., 2008) |
| *spsM::kan* (GM3248) | *spsM::kan\** | (Sanchez-Vizuet et al., 2015) |
| *spsM::kan amyE::spsM* (GM3326) | *spsM::kan amyE::spsM\* (spec)* | (Sanchez-Vizuet et al., 2015) |
| *spsM::kan* (MP) | *spsM::kan* | This work |
| *spsM::kan amyE::spsM* (MP) | *spsM::kan amyE::spsM (spec)* | This work |
| *spsM::kan* (GM3248)<br>*amyE::spsM* (MP) | *spsM::kan\* amyE::spsM (spec)* | This work |
| Lys SPβ | *spsM::SPβ* | This work |
| *spsM::kan* (GM3248)<br>*amyE::comP* | *spsM::kan\* amyE::P<sub>hs</sub>-comP (spec)* | This work |
| *spsM::kan* (GM3248)<br>*comP::ery* | *spsM::kan\* comP::ery* | This work |
| *spsM::kan amyE::spsM* (GM3326) *comP::ery* | *spsM::kan amyE::spsM\* comP::ery* | This work |
| *spsM::kan* (MP) *comP::ery* | *spsM::kan comP::ery* | This work |
| *spsM::kan amyE::spsM* (MP) *comP::ery* | *spsM::kan amyE::spsM comP::ery* | This work |
| ***E. coli*** |  |  |
| AEC1002 | pDR111::P<sub>hs</sub>-*comP* (*amp*, *spec*) | (Bareia et al., 2018) |
| DH5α pIC679 | pIC679 (*amp*, *spec*) | This work |
| MC1061 | phy-sGFP (*cm*) | (Dragoš, Andersen, et al., 2021) |
| MC1061 | pTB498 (*spec*) | (Dragoš, Kiesewalter, et al., 2018) |

\*The genotype listed is as described in the research article by Sanchez-Vizuet et al. (2015).

**Table 2:**
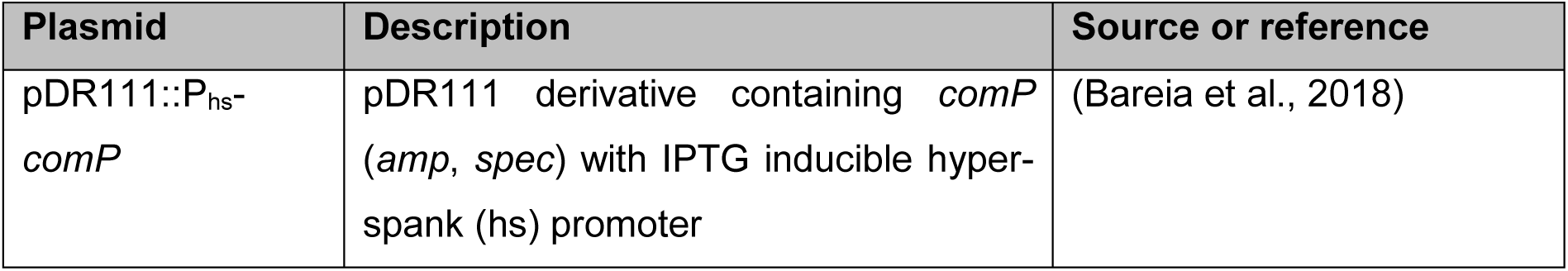
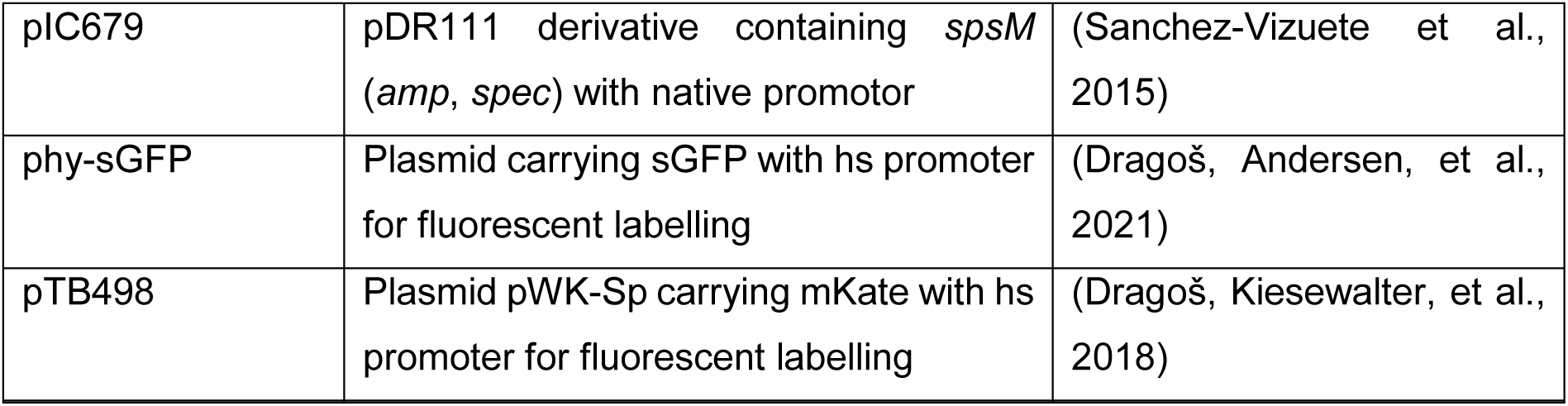
Plasmids used for *B. subtilis* mutant construction.

**Table 3:**
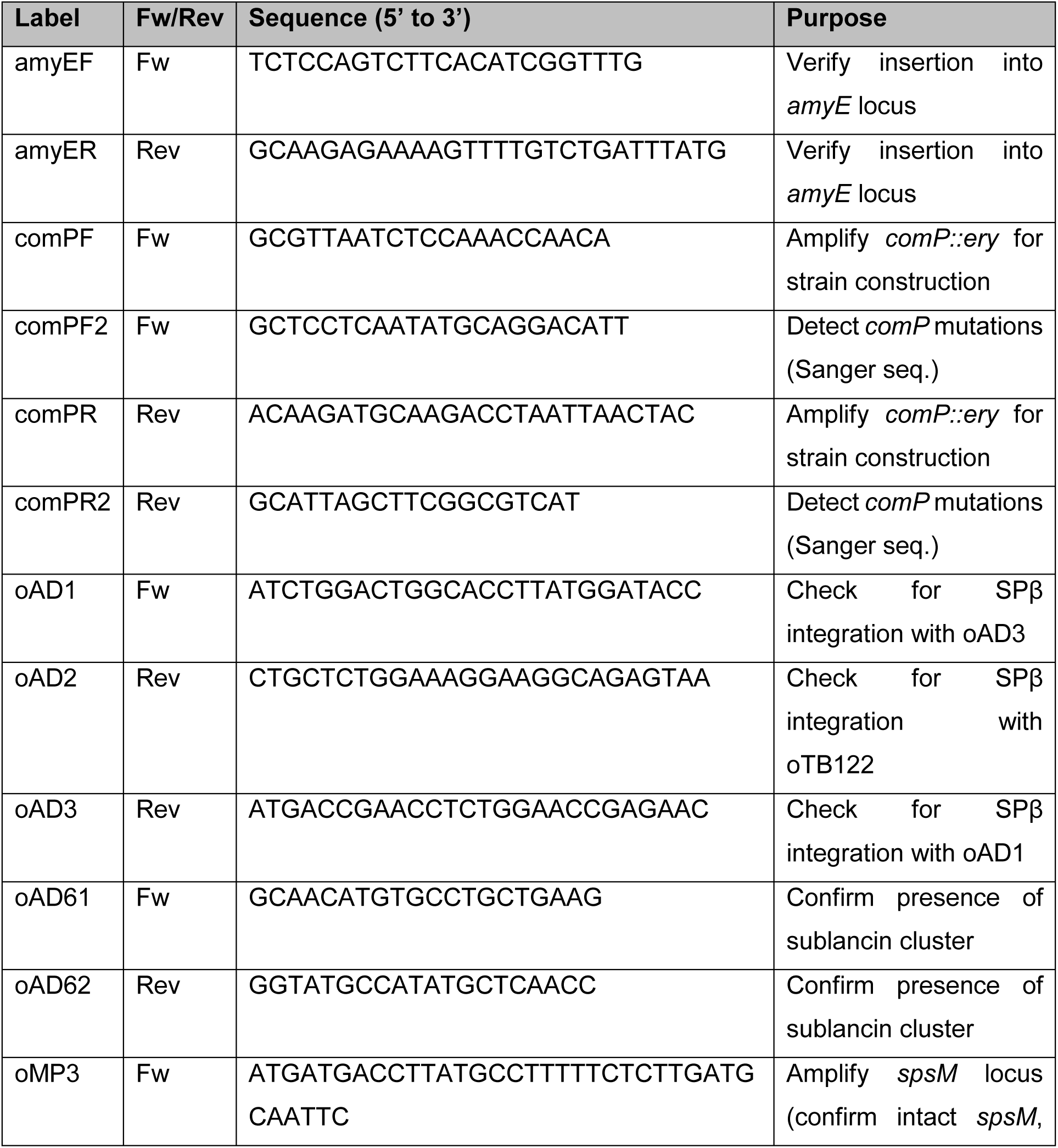
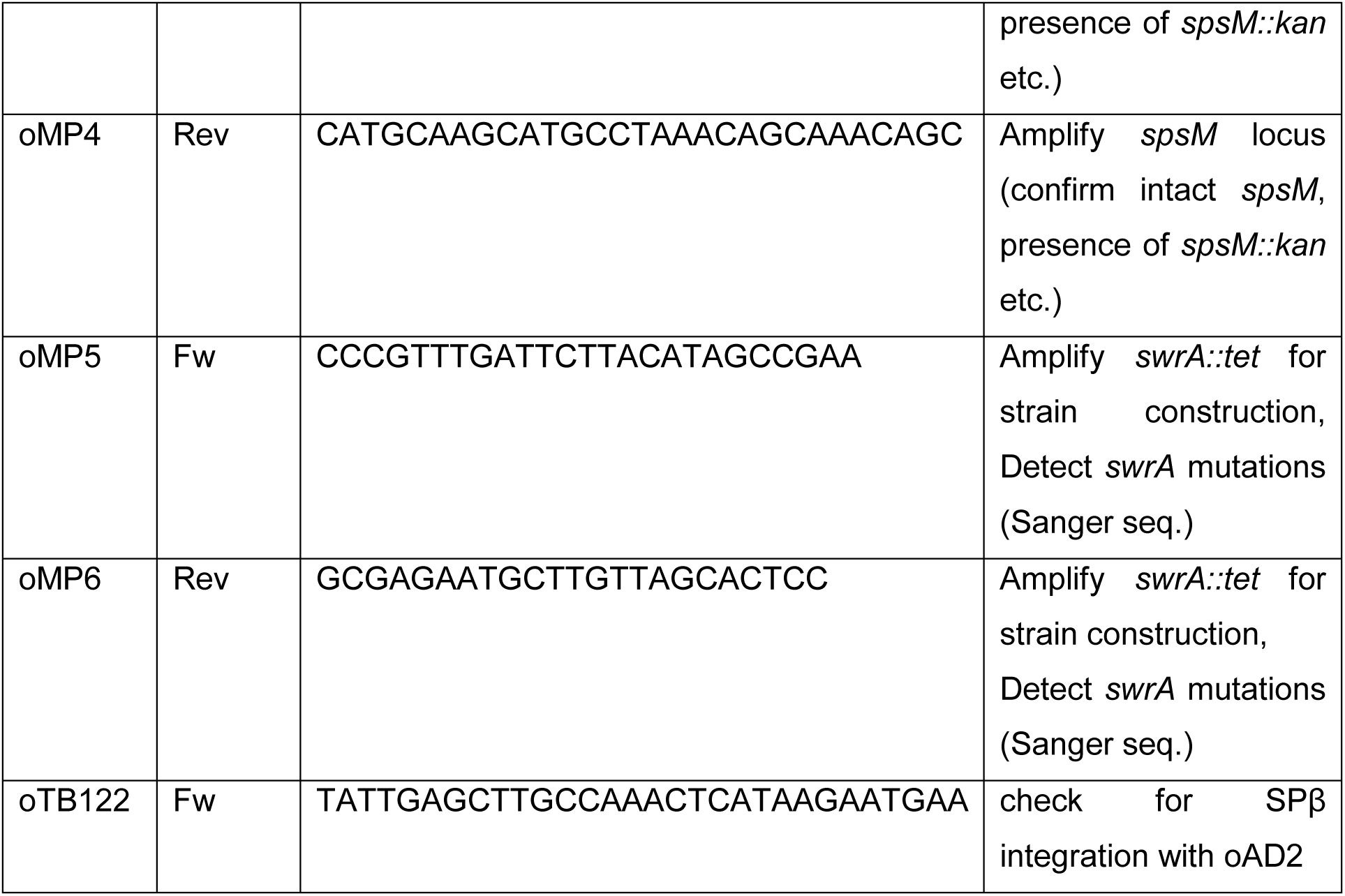
Oligonucleotides used in this work. Forward primers (Fw), Reverse primers (Rev).

**Table 4:** Strains analysed with whole-genome sequencing and their short read archive accession numbers.

| Strain | Accession number | Sequencing method |
| --- | --- | --- |
| P9_B1 *spsM::kan* | SRR31313896 | Illumina |
| P9_B1 fluffy | SRR31346685 | Illumina |
| NDmed wt | Raw reads: SRR32479380<br>Assembled(annotated):<br>CP183238 | PacBio |
| NDmed *spsM::kan* (GM3248) | SRR32463897 | Illumina |
| NDmed *spsM::kan amyE::spsM* (GM3326) | SRR31379056 | Illumina |
| NDmed *spsM::kan* (MP) | SRR32465634 | Illumina |
| NDmed Lys SPβ | SRR32478187 | Illumina |

Mutant *B. subtilis* strains prepared in this study were constructed by transforming the desired strains with polymerase chain reaction (PCR) products or with plasmids carrying the desired region (Danevčič et al., 2021). Successfully transformed mutants were obtained by selecting for desired antibiotic resistance. Working concentrations of antibiotics were 5 μg/ml kanamycin (Kan), 100 μg/ml spectinomycin (Spec), 5 μg/ml chloramphenicol (Cm), 10 μg/ml tetracycline (Tet) and 100 μg/ml ampicillin (Amp).

The NDmed *spsM::kan* (MP) and PS-216 *spsM::kan* strains were generated by transforming the NDmed wt and PS-216 wt strains with PCR product of the *spsM::kan* region from NDmed *spsM::kan* (GM3248) strain. The pIC679 plasmid (Table 2), carrying the *spsM* gene with its native promotor, was previously constructed and described in Sanchez-Vizuete et al. (2015). The pIC679 plasmid was transformed into *E. coli* DH5α strain (Danevčič et al., 2021; Sanchez-Vizuete et al., 2015) and was then isolated and used to generate strains prepared in this study labelled as *amyE::spsM*. Strains labelled as *swrA::tet* or *comP::ery* were constructed by transforming the desired strains with PCR product of the *swrA::tet* or *comP::ery* region from DS215 or BKE31690 strain, respectively. The pDR111::P_hs_-*comP* plasmid (Table 2) was isolated from strain AEC1002 and was used to construct the NDmed *spsM::kan* (GM3248) *amyE::comP* strain.

Fluorescently labelled strains were constructed by transforming the desired strains with the phy-sGFP or pTB498 plasmids (Table 2) that carry the sGFP and mKate fluorescent marker fused with the hyper-spank promotor, respectively. The NDmed wt, *spsM::kan* (GM3248) and *spsM::kan amyE::spsM* (GM3326) strains were previously constructed and described in Sanchez-Vizuete et al. (2015). The P9_B1 fluffy strain was obtained as a result of a spontaneous mutation in P9_B1 *spsM::kan* strain grown as macrocolonies on LB agar plates after 48 hours of incubation at 30°C.

The NDmed Lys SPβ strain was constructed by preparing a SPβ phage suspension from MB8_B7 strain, by prophage induction using 0.5 μg/ml mitomycin C. The phage suspension was then used to lysogenize the NDmed wt strain. The presence of SPβ phage in *spsM* gene was confirmed by PCR using the primer pairs presented in Table 3. The presence of the phage in the genome was additionally confirmed by Illumina whole-genome sequencing (Table 4). NDmed Lys SPβ was additionally verified through prophage activity test, by 0.5 μg/ml mitomycin C induction, followed by plaque assay, as previously described (Dragoš, Andersen, et al., 2021). Lysogeny was additionally confirmed with sublancin activity assay, as previously described (J.-Y. F. Dubois et al., 2009).

### Standardized plate preparation

A standardized protocol of solid media preparation was developed to ensure reproducibility of colony morphology across replicates. The medium was always prepared one day prior to inoculation using 250 mL Erlenmeyer flasks containing either 100 mL or 200 mL of medium with desired amount of agar. Following autoclaving, the medium was cooled to 55°C for 30 minutes (100 mL) or 45 minutes (200 mL), and 20 mL aliquots were then transferred into 90 mm Petri dishes. To ensure similar humidity for all plates they were left at room temperature to dry overnight without stacking. Immediately before inoculation, plates were dried in a laminar flow hood for a specific duration tailored to each experimental protocol.

### Macrocolony experimental conditions and image analysis

To compare macrocolony morphologies among *B. subtilis* mutant strains, cultures were grown on various solid media containing 1.5% agar that were prepared using the standardized plate preparation protocol described above. Colony morphology was assessed on LB agar (Condalab, Spain), TSA (Condalab, Spain), and LBGM agar, which enhances extracellular matrix production in *B. subtilis* biofilm (Gingichashvili et al., 2020; Shemesh & Chai, 2013; Werb et al., 2017). LBGM medium was prepared by supplementing LB agar with sterile 1% glycerol and 100 μM MnSO₄ post-autoclaving. For strains harbouring constructs under the control of hyper-spank promoter, the media was supplemented with 0.1 mM IPTG. To initiate macrocolony growth 10 μL of overnight culture was spotted onto the plates, dried for 5 minutes post-inoculation and incubated at 30°C for 48 hours. To ensure consistency in macrocolony morphology across conditions, different strains were grown on the same plates, and each condition was tested at least in duplicate.

Macrocolony images were acquired using Leica MZ FLIII and Nikon SMZ25 stereomicroscopes at the infrastructural center “Microscopy of Biological Samples” (Biotechnical Faculty, University of Ljubljana, Slovenia). Surface structure of macrocolony biofilms was analysed with SurfCharJ plugin (Chinga et al., 2007) in Fiji (ImageJ) (Schindelin et al., 2012). Pixel intensity in the 2D images was assumed to correlate with object height, with brighter regions representing more protruding surface features. The colony morphology was analysed by pairwise correlation analysis implemented in custom C++ program routine (Dogsa et al., 2023). First, by using Fiji (ImageJ), the RGB image was split into three colour channels. For further analysis we have always chosen the green channel image, where background of the colony was subtracted. Then, all non-background pixel intensity values were exported, together with corresponding positional coordinates. These served as input data to a C++ program routine, which calculates the normalized pairwise correlation function defined as:

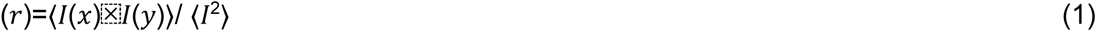

where < I(x)I(y)> is the averaged product of fluorescence intensities of two pixels at a distance r and <I^2^ > is the averaged square of intensities of all pixels, making γ(0) = 1. In images containing larger objects, the pairwise correlation function decays more slowly with distance. Oscillations in the function reflect repeating spatial motifs and local ordering. In contrast, images composed of randomized, unstructured pixels, exhibit a rapid decay of the correlation function toward zero, with no undulations.

Macrocolony morphology of original stocks of the NDmed *spsM::kan* mutants (Sanchez-Vizuete et al., 2015) GM3248 clone 1 and GM3248 clone 2 was tested by the research team at the Micalis Institute. To perform the macrocolony assay, 6-well plates were prepared by adding 4 mL of LB (BD Difco™, USA) with 1,5% agar or TSA (BioMérieux, France) with 1,5% agar to each well. The plates were dried under a laminar flow hood for 1.5 hours and then left at room temperature for 5 days prior to use. For inoculum preparation, a single colony was grown overnight in 5 mL of LB or TSB, supplemented with kanamycin (8 µg/mL) for the mutant strains, at 30 °C with shaking at 180 rpm for 18 hours. Cultures were centrifuged at 5000 × g for 5 minutes and resuspended in fresh medium. A 4 µL aliquot of the resuspension was spotted at the center of each well, and plates were incubated at 30 °C to allow macrocolony development.

### Swimming and swarming motility assays

Swimming and swarming motility of *B. subtilis* strains was assessed on plates prepared using the standardized plate preparation protocol described above. Swimming motility was tested on LB soft agar plates containing 0.3% agar, while swarming motility was assessed on LB semi-solid agar plates with 0.7% agar, using an adjusted version of a previously described protocol (Nordgaard et al., 2022). For strains harboring constructs under the hyper-spank promoter, the media was supplemented with 0.1 mM IPTG. Plates were prepared one day prior to use and dried overnight at room temperature without stacking. Immediately before inoculation, 0.3% agar plates were dried in a laminar flow hood for 5 minutes, and 0.7% agar plates were dried for 20 minutes to standardize surface moisture. 2 μL of culture was spotted at the center of the prepared plates, which were then incubated at 37°C for up to 8 h without stacking, to maintain uniform humidity. Motility was quantified as the mean colony diameter measured along two perpendicular axes.

### Competition assay

The competition assay was conducted, using standardized macrocolony preparation as described above. Strain mixtures were grown on LB and LBGM solid media (1.5% agar) prepared using the standardized plate preparation protocol described above. Overnight cultures of *B. subtilis* P9_B1 strains were adjusted to equal optical density at 600 nm (OD₆₀₀). Strain pairs, each tagged with either GFP or mKate fluorescent reporters, were mixed in equal volumes (100 μL each; 1:1 v/v). From each mixture, 10 μL was spotted onto LB or LBGM agar plates and incubated at 30 °C for 48 h. To correct for potential differences in reporter fluorescence intensity, monocultures of each fluorescent strain were grown in parallel on the same media.

Macrocolony images were acquired using a Nikon SMZ25 stereomicroscope at the infrastructural center “Microscopy of Biological Samples” (Biotechnical Faculty, University of Ljubljana, Slovenia). Fluorescence quantification was performed in Fiji (ImageJ) (Schindelin et al., 2012) by measuring the integrated density of the green (GFP) and red (mKate) channels. Values from cocultures were normalized against monoculture fluorescence intensities to account for potential variability in reporter expression. Relative abundance of each strain in coculture was expressed as a percentage of the total normalized fluorescence signal.

### Spontaneous mutation screening

To assess whether disruption of the *spsM* gene promotes spontaneous diversification and the emergence of mutations, we conducted a screening assay based on conditions under which such mutations were first observed. A volume of 10 µL from overnight cultures was spotted onto LB agar plates (Condalab, Spain), briefly dried for 5 minutes, and incubated at 30°C for 48 hours. All strains were plated on the same LB agar plate as separate macrocolonies to minimize variability in growth conditions.

Following incubation, peripheral outgrowths that developed at the edges of macrocolonies and displayed distinct morphology compared to the colony center were identified. Both peripheral and central regions were restreaked on fresh LB agar plates, and colony morphology was assessed after incubation. Outgrowths that consistently showed different morphology from the central colony were grouped by visual phenotype. A subset of these strains, representing each morphological group, was selected for Sanger sequencing of the *swrA* and *comP* loci using primer pairs oMP5/oMP6 and comPF2/comPR2 (Table 3). Within each group, strains exhibited identical sequencing results, allowing classification based on shared morphological features. One representative strain from each group was chosen for further phenotypic analysis, including assessment of macrocolony morphology and swimming and swarming motility performed as described above.

The assay was repeated across four independent experiments. Minor differences in environmental conditions, such as humidity and LB agar batch, may have influenced the frequency of variant emergence. Therefore, results from each experiment are presented independently. We did not perform statistical comparisons or aggregate the data across replicates, as the purpose of this analysis was to qualitatively document the consistent association between *spsM* disruption and spontaneous mutation emergence.

### Whole genome sequencing and sanger sequencing

Whole genome sequencing (WGS) of *B. subtilis* P9_B1 and NDmed mutant strains was performed by SeqOmics Biotechnology Ltd. using an Illumina NextSeq platform. Paired-end libraries (2 × 150 bp, ∼10 million reads per sample, ∼700× coverage) were prepared with the NEBNext® Ultra™ II DNA Library Prep Kit and sequenced with the NextSeq 500/550 High Output Kit v2. Reads were mapped to the corresponding reference genome assemblies (NDmed: ASM74047v1 and ASM4853740v1; P9_B1: ASM966245v1), and single nucleotide polymorphism (SNP) analysis was conducted using CLC Genomics Workbench 23.0.4.

For NDmed strains, initial mapping used the draft reference genome assembly (ASM74047v1, 10 contigs). To improve resolution, a complete circular genome of NDmed wild-type was generated de novo using PacBio Sequel II (Novogene Europe), and annotated with Prokka (Seemann, 2014). High-quality Illumina reads were aligned to this reference using Bowtie (Langmead et al., 2009), and SNPs were identified using SAMtools (Li et al., 2009), following quality checks with MultiQC (Ewels et al., 2016). Detected SNPs in *swrA* and *comP* were validated via PCR and Sanger sequencing (Macrogen, Maastricht) using primer pairs oMP5/oMP6 and comPF2/comPR2 (Table 3).

### Bioinformatic and statistical analysis

Statistical analysis for all measurements was performed with ANOVA and Tukey’s Honest Significant Difference method (p < 0.05) in RStudio using aov() and TukeyHSD() functions, respectively. Function multcompLetters4() was used to obtain the compact letter display from a performed ANOVA. ExPASy translate tool (Gasteiger et al., 2003) was used to translate the DNA sequences acquired with Sanger sequencing to analyse their potential effect on translated protein sequences. All nucleotide sequences were viewed, analysed and aligned with the program Snapgene (GSL Biotech) by using the algorithm MUSCLE (MUltiple Sequence Comparison by Log-Expectation) (Edgar, 2004; Madeira et al., 2024).

## RESULTS

### Emergence of new morphotype observed in *B. subtilis* P9_B1 with disruption of prophage integration locus *spsM*

The primary objective of this study was to explore the eco-evolutionary consequences of prophage integration into the functional *B. subtilis* locus *spsM*. Previous research indicated that insertion of the SPβ prophage, which is present in common laboratory strains such as *B. subtilis* 168 and NCBI 3610, disrupts *spsM* and negatively affects biofilm formation. In contrast, the clinical isolate *B. subtilis* NDmed, which lacks the SPβ prophage and carries an intact *spsM* gene, was shown to form robust biofilms (Sanchez-Vizuete et al., 2015).

To evaluate the impact of the *spsM::kan* mutation on colony morphology, macrocolonies of the *B. subtilis* soil isolate P9_B1 were repeatedly grown on standardized solid LB medium. In this background, deletion of *spsM* did not produce the colony morphology changes previously reported (Sanchez-Vizuete et al., 2015) for the *B. subtilis* NDmed strain (Figure 1A). Specifically, it did not lead to the development of a flat phenotype, which is characterized by smooth, unstructured colonies lacking the typical wrinkled architecture. In contrast, we were able to replicate the characteristic flat colony morphology reported by Sanchez-Vizuete et al. (2015) using the NDmed *spsM::kan* (GM3248) strain (Figure 1A). However, when we lysogenized NDmed and P9_B1 with the SPβ prophage, we did not observe the expected flat colony phenotype (Figure 1A), even though molecular validation and whole-genome sequencing confirmed that SPβ had integrated at the *spsM* locus. Confirmation of lysogenization for NDmed is presented in Supplementary Figure S1, while the lysogenic state of P9_B1 has been previously verified (Dragoš, Andersen, et al., 2021).

**Figure 1:**
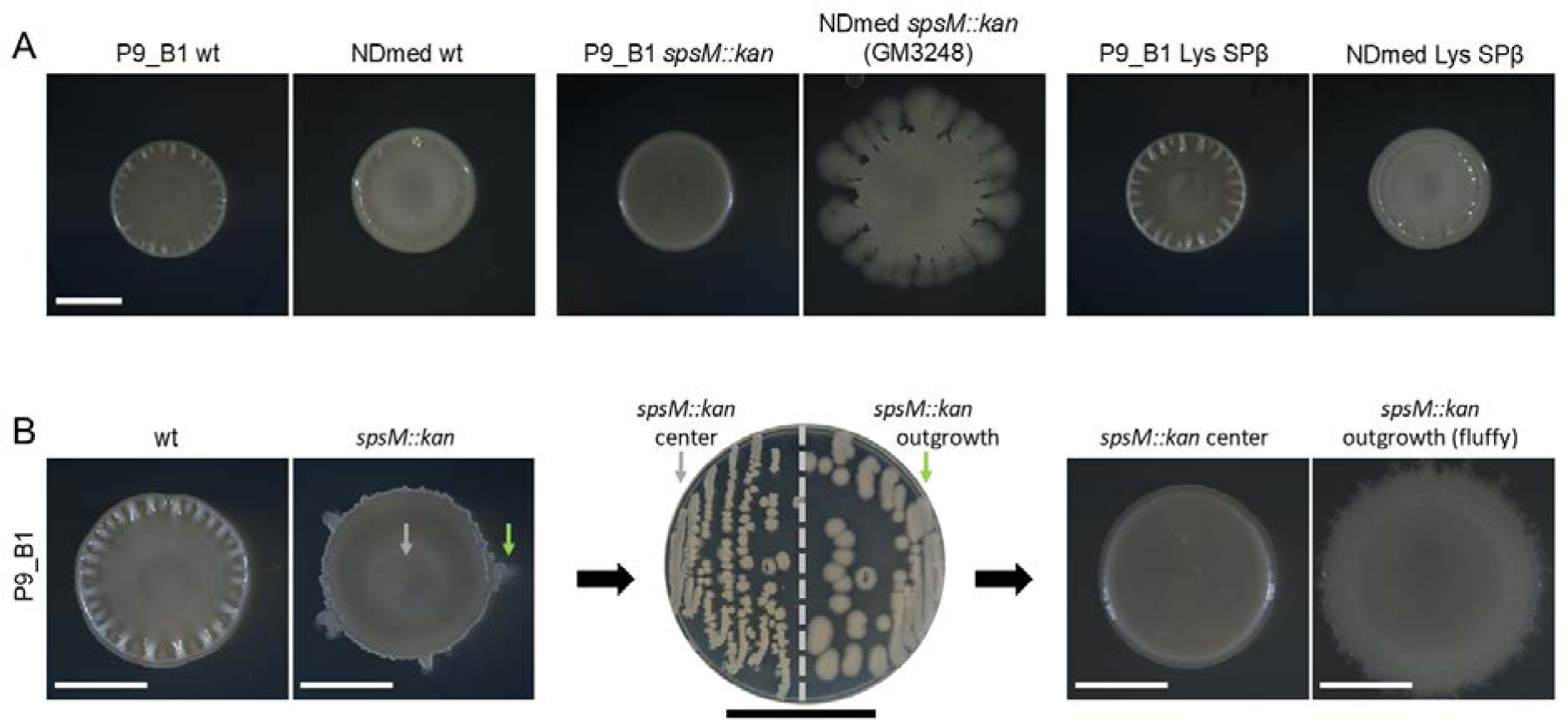
Morphological comparison of *B. subtilis* macrocolonies grown on standardized solid LB medium. **A** Macrocolony morphology of P9_B1 and NDmed strains grown on LB agar at 30 °C for 48 hours. Scale bar: 5 mm. **B** Macrocolony morphology of the P9_B1 *spsM::kan* strain and its derivatives. All strain labels refer to P9_B1 strains. On the left, macrocolonies of the wild-type (wt) and *spsM::kan* mutant are shown after 48 hours of growth on LB agar at 30 °C. The middle panel shows an LB plate with isolated colonies derived from streaking the center (grey arrow) and outgrowth (green arrow) regions of the *spsM::kan* macrocolony. On the right are macrocolonies formed by the isolated strains: the *spsM::kan* center and the outgrowth strain referred to as “fluffy”. White scale bar: 5 mm. Black scale bar: 5 cm.

Interestingly, on solid LB medium, instead of displaying the expected morphology changes, the P9_B1 *spsM::kan* strain occasionally developed peripheral outgrowths that were absent in the wild-type strain (Figure 1B), suggesting spontaneous diversification through the appearance of mutations. These outgrowths typically appeared after 48 hours of incubation at 30 °C. To determine whether the outgrowths resulted from a spontaneous mutation, we transferred both the center and the outgrowth of a single macrocolony onto fresh solid LB medium and examined the resulting colony morphologies (Figure 1B). The macrocolonies derived from the outgrowth displayed a distinct morphology compared to both the original P9_B1 *spsM::kan* strain and the central region of the same macrocolony. This newly emerged lineage of *spsM::kan* was named “fluffy”, due to an irregular colony edge resembling a cotton wool. The “fluffy” morphotype was stable and could be recreated from −80°C stock, followed by overnight culture and colony spotting, indicating it was caused by an additional mutation that emerged in the *spsM::kan* strain background (Figure 1B).

### ‘Fluffy’ morphotype is caused by spontaneous mutation in *swrA* which rapidly arises in *spsM::kan* genetic background

Illumina whole-genome resequencing was performed on genomic DNA of both the ancestral P9_B1 *spsM::kan* strain (NCBI SRA accessions: SRR31313896) and the morphologically distinct spontaneously derived mutant strain, hereafter referred to as P9_B1 fluffy (NCBI SRA accessions: SRR31346685). A single mutation unique to the P9_B1 fluffy strain was identified: a thymine deletion at position 26 of the *swrA* coding sequence (swrA:c.26delT), resulting in a frameshift mutation (Figure 2). These findings indicate that the P9_B1 *spsM::kan* strain went through a fast spontaneous genetic diversification within 48 hours of growth on solid LB medium at 30 °C.

**Figure 2:**
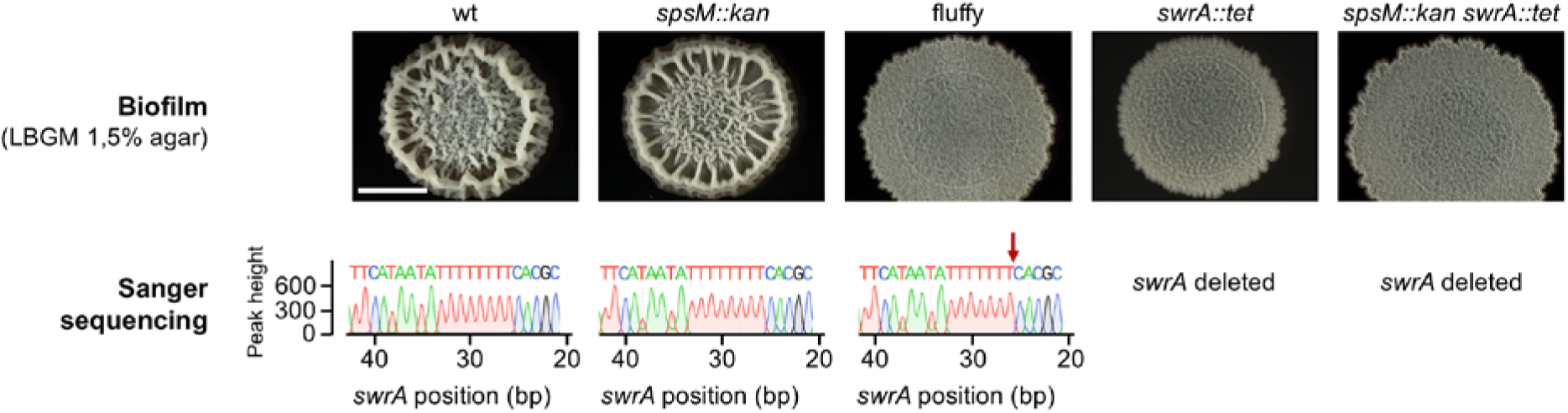
Biofilm morphology and Sanger sequencing confirmation of a spontaneous *swrA* mutation in the P9_B1 fluffy strain. All strain labels refer to P9_B1 derivatives. The “fluffy” strain refers to the *spsM::kan* strain carrying the spontaneous *swrA* c.26delT mutation. The top panel shows macrocolony biofilms of strains with (wt and *spsM::kan*) and without (fluffy, *swrA::tet*, *spsM::kan swrA::tet*) an active *swrA*. Macrocolony biofilms were grown on solid LBGM medium and imaged after 48 h incubation at 30°C. Scale bar: 5 mm. The bottom panel displays segments of Sanger sequencing results covering positions 20–40 bp of the *swrA* gene. The nucleotide sequences derived from Sanger reads are shown for each strain, with the corresponding peak height plotted on the Y-axis. The spontaneous swrA:c.26delT mutation in the fluffy strain is indicated by a red arrow and corresponds to a frameshift mutation not observed in the other strains.

To assess whether the spontaneous *swrA* mutation in the P9_B1 fluffy strain affects macrocolony biofilm morphology, we examined macrocolonies on solid LBGM medium (Figure 2), which promotes biofilm formation by enhancing extracellular matrix production (Gingichashvili et al., 2020; Shemesh & Chai, 2013; Werb et al., 2017). The morphology of P9_B1 fluffy was considerably different from that of its ancestral P9_B1 *spsM::kan* strain. Macrocolony biofilm of P9_B1 fluffy was more widely spread, flatter, and exhibited smaller wrinkles (Figure 2). To confirm these changes in morphology resulted from the *swrA* frameshift mutation we constructed *swrA* disruption mutants P9_B1 *swrA::tet* and P9_B1 *spsM::kan swrA::tet*. P9_B1 fluffy and P9_B1 *spsM::kan swrA::tet* mutants exhibited indistinguishable morphologies, supporting the conclusion that the *swrA* mutation leads to altered macrocolony biofilm morphology in P9_B1 fluffy.

### Spontaneous mutation in *swrA* in P9_B1 fluffy strain leads to loss of swarming and impaired swimming motility

In *B. subtilis*, SwrA regulates both swimming and swarming by promoting flagellar biosynthesis (Mukherjee & Kearns, 2014). Since the P9_B1 fluffy strain resulted from a spontaneous mutation in *swrA*, we compared motility of wild-type and mutant strains on LB medium containing 0.3% (swimming) and 0.7% (swarming) agar (Figure 3). P9_B1 wt and *spsM::kan* strains displayed similar swimming and swarming motility patterns. During swimming, both strains formed uniform, featureless colonies, whereas under swarming conditions, they developed structured patterns with compact zones and dendritic extensions. In contrast, all three strains carrying the *swrA* mutation were significantly impaired in both modes of motility. Swimming motility was still present in *swrA* mutants but was statistically significantly lower than in strains with intact *swrA* (Figure 3B, 3D). Notably, the swimming patterns of P9_B1 fluffy and *spsM::kan swrA::tet* strains were inconsistent across replicates, sometimes displaying thin dendrites and other times not. In contrast, all *swrA* mutant strains completely lost the ability to swarm (Figure 3B, 3C). These results confirm that the spontaneous *swrA* frameshift mutation in P9_B1 fluffy leads to a loss of swarming motility and altered macrocolony morphology.

**Figure 3:**
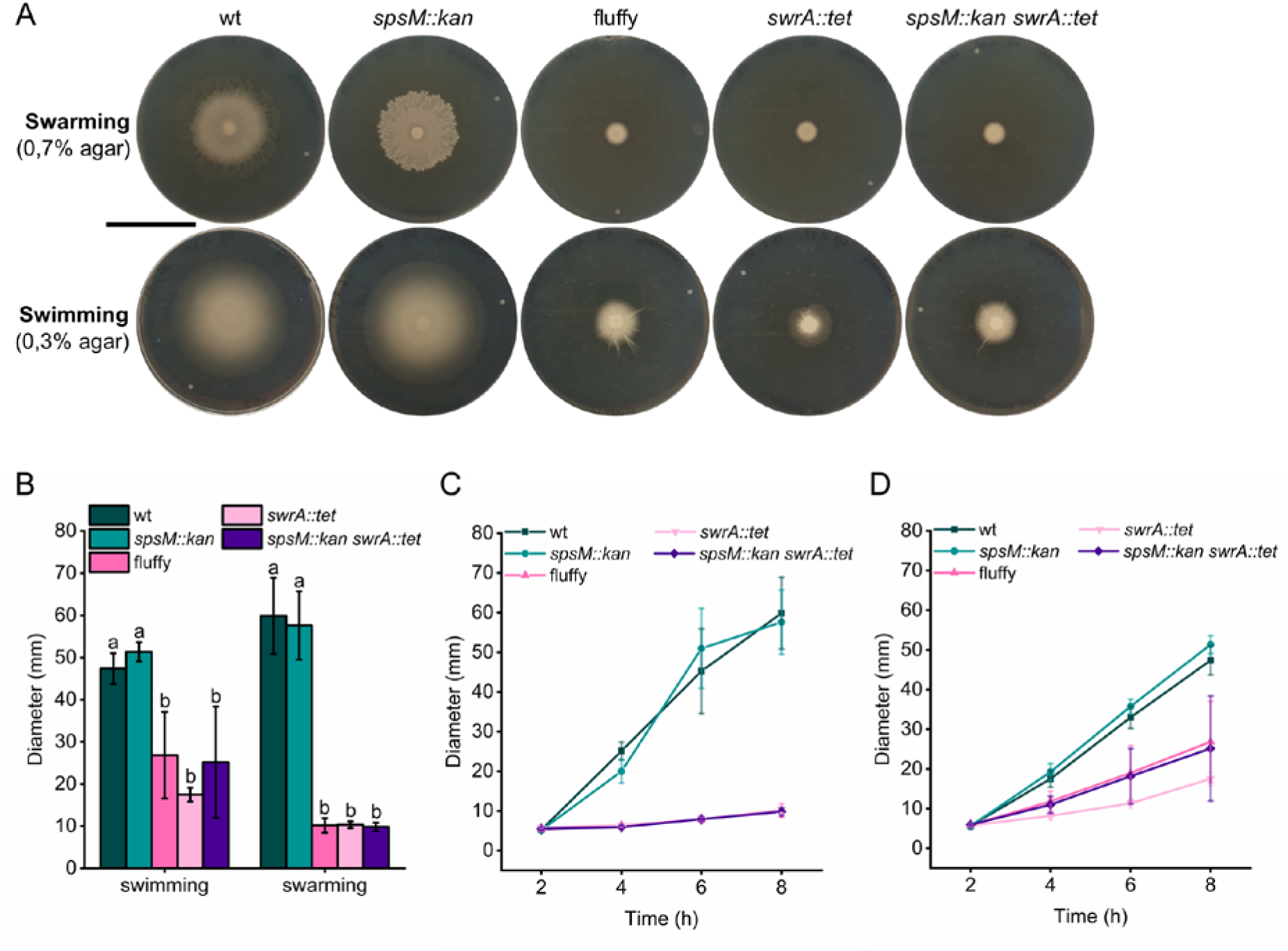
Decreased swimming and loss of swarming motility resulting from a spontaneous mutation of the *swrA* gene in the P9_B1 fluffy strain. All strain labels refer to P9_B1 derivatives. The fluffy strain refers to the *spsM::kan* strain carrying the spontaneous *swrA* c.26delT mutation. **A** Images of swarming and swimming motility of strains with (wt and *spsM::kan*) and without (fluffy, *swrA::tet*, *spsM::kan swrA::tet*) an active *swrA*. Swimming assays were performed on LB with 0.3% agar, and swarming assays on LB with 0.7% agar. Swimming and swarming motility plates were imaged after 8 h incubation at 37°C. Plates were imaged against a black background so that bacterial colonization zones appear white, and uncolonized agar appears dark. Scale bar: 5 cm. **B** Quantification of swimming and swarming diameters after 8 hours at 37°C. Columns represent mean diameters (n=6) and error bars represent standard deviation. Statistical analysis was performed using one-way ANOVA followed by Tukey’s Honest Significant Difference (HSD) test. Letters above error bars indicate statistically significant differences between strains separately for swimming and swarming (p < 0.05). **C, D** Dynamics of swarming (C) and swimming (D) motility over an 8-hour period, with measurements taken at 2 h, 4 h, 6 h, and 8 h.

### Flat morphotype in *B. subtilis* NDmed *spsM::kan* (GM3248) is associated with spontaneous mutation in *comP*

We further investigated why disruption of *spsM* (*spsM::kan*) leads to pronounced changes in biofilm morphology in the *B. subtilis* clinical isolate NDmed but not in the soil isolate P9_B1, and why we were unable to recreate the flat colony morphotype in SPβ lysogen of NDmed. In order to do that, we tested the effects of *spsM* complementation, we recreated *spsM* deletion in NDmed strain and we reexamined macrocolony biofilms on other media types like TSA, and LBGM (Figure 4). NDmed strains labeled as wt, *spsM::kan* (GM3248) and *spsM::kan amyE::spsM* (GM3326) were previously described by Sanchez-Vizuete et al. (2015), while strains labeled with (MP) were constructed as part of this study to independently verify the phenotypic effects of the *spsM::kan* mutation in the NDmed background.

**Figure 4:**
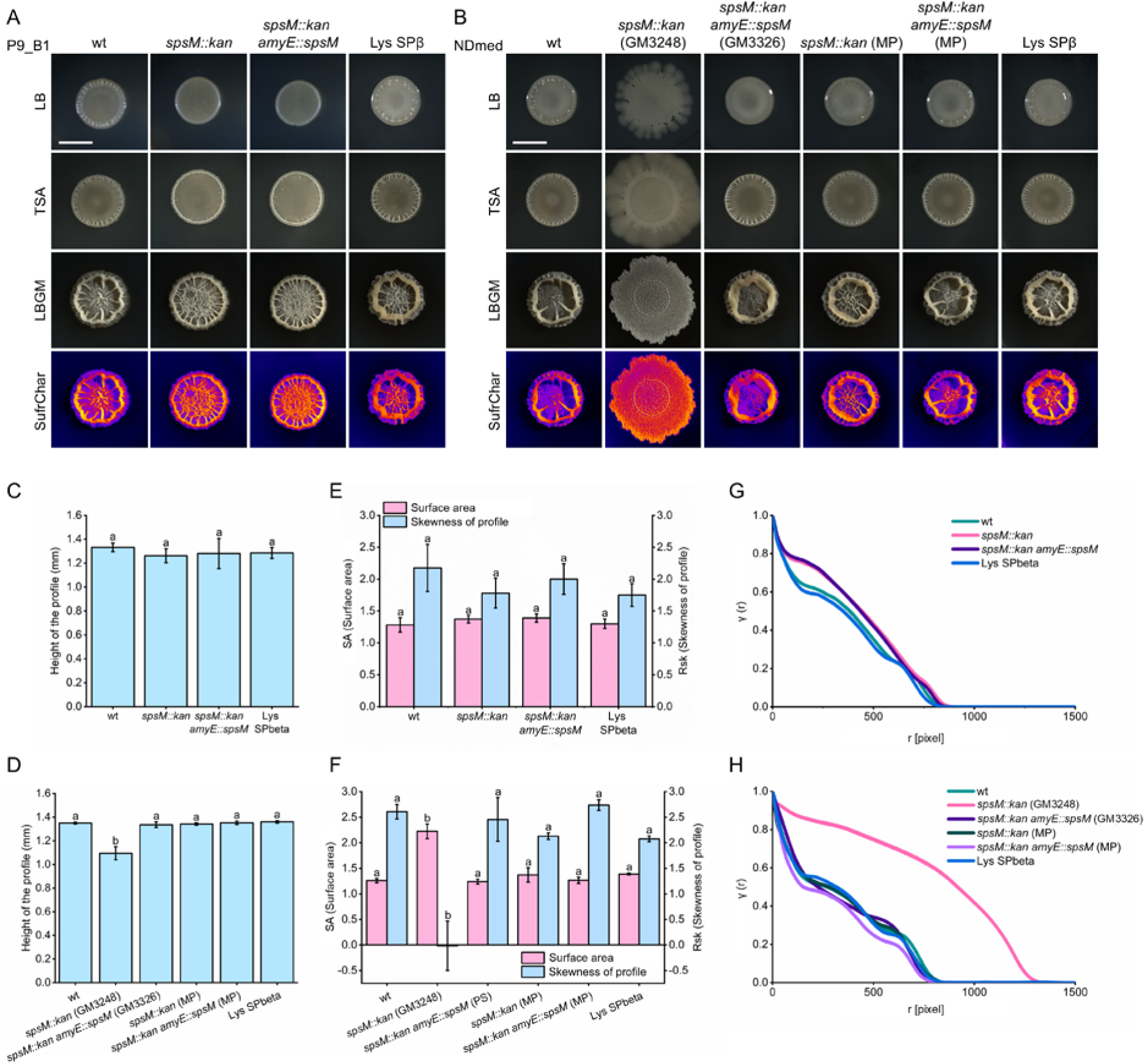
Macrocolony biofilm morphology, surface characterization, and pairwise correlation analysis of *B. subtilis* P9_B1 and NDmed strains. The NDmed strains labeled as wt, *spsM::kan* (GM3248), and *spsM::kan amyE::spsM* (GM3326) were kindly provided by Sanchez-Vizuete *et al*. (2015). Strains marked with (MP) were constructed in this study to independently confirm the phenotypic effects of the *spsM::kan* mutation in the NDmed background. **A, B** Macrocolony biofilm morphology of P9_B1 strains (A) and NDmed strains (B) grown on LB, TSA, and LBGM media at 30°C for 48 hours. For LBGM-grown biofilms, surface characteristics were further analyzed using the SurfCharJ plugin in Fiji (ImageJ). Scale bar: 5 mm. **C–F** Quantitative analysis of biofilm structure for P9_B1 (**C, E**) and NDmed (**D, F**) strains based on SurfCharJ analysis: **C, D** Height of the biofilm profile; **E, F** surface area in relative units and skewness of the biofilm profile. Columns represent mean values (P9_B1: n=6; NDmed: n=4) and error bars indicate standard deviation. Statistical analysis was performed using one-way ANOVA followed by Tukey’s Honest Significant Difference (HSD) test. Letters above error bars denote statistically significant differences between strains (p < 0.05). **G, H** Pairwise correlation analysis of macrocolony biofilms grown on LBGM media, performed for P9_B1 (**G**) and NDmed (**H**) strains.

As expected, based on our initial results on LB, we were again successfully able to reproduce previous findings on TSA and LBGM, where NDmed *spsM::kan* (GM3248) strain formed wider, flatter macrocolonies compared to the NDmed wt strain, which formed smaller macrocolonies with prominent large wrinkles (Figure 4B). In addition, the NDmed *spsM::kan amyE::spsM* (GM3326) strain showed successful complementation, displaying a macrocolony morphology identical to that of the NDmed wt strain, consistent with the results reported by Sanchez-Vizuete et al. (2015). In contrast, the NDmed mutant strains we constructed for this study (labeled MP) behaved differently. The NDmed *spsM::kan* (MP) strain exhibited no major morphological differences from the NDmed wt strain, nor did the complemented NDmed *spsM::kan amyE::spsM* (MP). We also constructed a *spsM::kan* mutant of *B. subtilis* soil isolate PS-216 and observed no effect on biofilm morphology (Supplementary Figure S2A). Additionally, pellicle biofilms were examined across all strains, but no major differences in biofilm morphology were detected (Supplementary Figure S2B).

To quantitatively assess whether the *spsM::kan* mutation affects biofilm morphology, we characterized the surface structure and spatial segregation of P9_B1 and NDmed macrocolony biofilms grown on LBGM medium. Among all the strains tested, only the NDmed *spsM::kan* (GM3248) mutant showed statistically significant differences in surface features, specifically in maximum profile height (Rt), surface area in relative units (SA), and profile skewness (Rsk) (Figure 4C–F). Compared to the other NDmed and P9_B1 strains, which formed smaller, highly wrinkled macrocolonies, the NDmed *spsM::kan* (GM3248) mutant produced wider and flatter macrocolonies with a lower maximum height (Rt = 1.095 mm vs. ∼1.3 mm), increased surface area (SA = 2.2 vs. ∼1.3), and markedly reduced skewness (Rsk = –0.015 vs. 2.0–2.7). These measurements suggest a shift from prominent large-scale wrinkling to a higher density distribution of fine-scale surface structures. In contrast, the NDmed and P9_B1 strain *spsM::kan* mutants constructed for this study showed only minor effects on Rsk, none of which were statistically significant compared to their respective wild-type strains. A similar trend was observed in pairwise correlation analyses. Only the NDmed *spsM::kan* (GM3248) strain displayed a noticeably different pairwise correlation function compared to all other spsM::kan mutants (Figure 4G and 4H). Its correlation function is almost devoid of any oscillations that are present in other strains, suggesting significantly different structural order. Together, the surface characterization and pairwise correlation analysis in Figure 4 suggest that disruption of *spsM*, whether by antibiotic cassette or prophage integration, does not alter the biofilm phenotype of *B. subtilis*. These findings differ from earlier reports of a significant role for *spsM* in biofilm morphology in the NDmed strain (Sanchez-Vizuete et al., 2015).

A further interesting observation was that the biofilm morphology of NDmed *spsM::kan* (GM3248) strain (Figure 4) was similar to that of the spontaneously mutated P9_B1 fluffy strain (Figure 2). Both strains show flatter, more widely spread macrocolony biofilm with smaller more densely distributed wrinkles. This similarity led us to suspect that the NDmed *spsM::kan* (GM3248) strain might carry an additional spontaneous mutation contributing to the altered morphology. To investigate this, the NDmed strains were sent for whole-genome sequencing (NCBI SRA accessions: SRR31379056, SRR32463897, SRR32465634, SRR32478187, SRR32479380). *De novo* sequencing and genome assembly of the NDmed wt strain was also performed using PacBio technology (GenBank accession: CP183238). This assembly was used as a reference to identify mutations in related strains. Among the detected variants, only one mutation was unique to the NDmed *spsM::kan* (GM3248) strain and was not present in any of the other NDmed derivatives. This mutation was a thymine-to-adenine substitution at position 1222 of the *comP* coding sequence (comP:c.1222T>A). Bioinformatic analysis predicted that this substitution introduces a premature stop codon at the point of the mutation, likely resulting in a truncated, nonfunctional ComP protein. Notably, a secondary start codon located 15 bp downstream may allow for expression of the C-terminal portion of ComP as a separate polypeptide. Together, these results led us to suspect that the altered macrocolony biofilm morphology observed in the NDmed *spsM::kan* (GM3248) strain might be due to a spontaneous *comP* mutation, rather than the *spsM* disruption alone.

Following this observation, two original stocks of the NDmed *spsM::kan* mutant (Sanchez-Vizuete et al., 2015), labeled GM3248 clone 1 and GM3248 clone 2, were tested for altered macrocolony morphology and screened for presence of *comP* mutation by the research team at the Micalis Institute. The GM3248 clone 2 strain displayed the altered macrocolony morphology previously described by Sanchez-Vizuete et al. (2015) and carried the same *comP* mutation as detected in the NDmed *spsM::kan* (GM3248) copy of the strain stored at Biotechnical Faculty, University of Ljubljana (Supplementary Figure 3). These findings demonstrate that the morphological phenotype originally attributed directly to *spsM* disruption in NDmed is likely linked to an undetected spontaneous mutation in *comP*, and that disruption of *spsM* alone does not substantially alter biofilm morphology in either NDmed, P9_B1 or PS-216 genetic backgrounds.

### Complete deletion and spontaneous mutation of *comP* lead to altered colony biofilm morphology and impaired swarming motility

To determine whether the spontaneous *comP* mutation was responsible for the altered macrocolony biofilm morphology observed in the NDmed *spsM::kan* (GM3248) strain, we first constructed a complemented strain, NDmed *spsM::kan* (GM3248) *amyE::spsM* (MP). As expected, restoring *spsM* did not revert the morphology to wild-type (Figure 5A), supporting our hypothesis that the phenotype was not caused by disruption of *spsM*. We then introduced a copy of *comP* under IPTG-inducible control into the same strain (NDmed *spsM::kan* (GM3248) *amyE::comP*). Upon IPTG induction, the strain recovered wild-type morphology, confirming that the spontaneous *comP* mutation was indeed responsible for the altered biofilm phenotype. To test whether the premature stop codon in *comP* (comP:c.1222T>A) has the same effect as a complete gene deletion, we introduced *comP::ery* deletion in both NDmed *spsM::kan* (MP) and NDmed *spsM::kan* (GM3248). Both deletion strains showed identical macrocolony morphologies, which differed from their respective parental strains (Figure 5A). This further supports that the NDmed *spsM::kan* (GM3248) strain carries a nonfunctional *comP* allele due to the spontaneous mutation. Other strains with *comP::ery* mutation were also constructed and tested on different types of media and all had the same morphology, which was different from the *comP* spontaneous mutation comP:c.1222T>A (Supplementary Figure S4). Together, these results indicate that the premature stop codon in *comP* alters macrocolony biofilm morphology, but its effect is distinct from that of a complete *comP* deletion.

**Figure 5:**
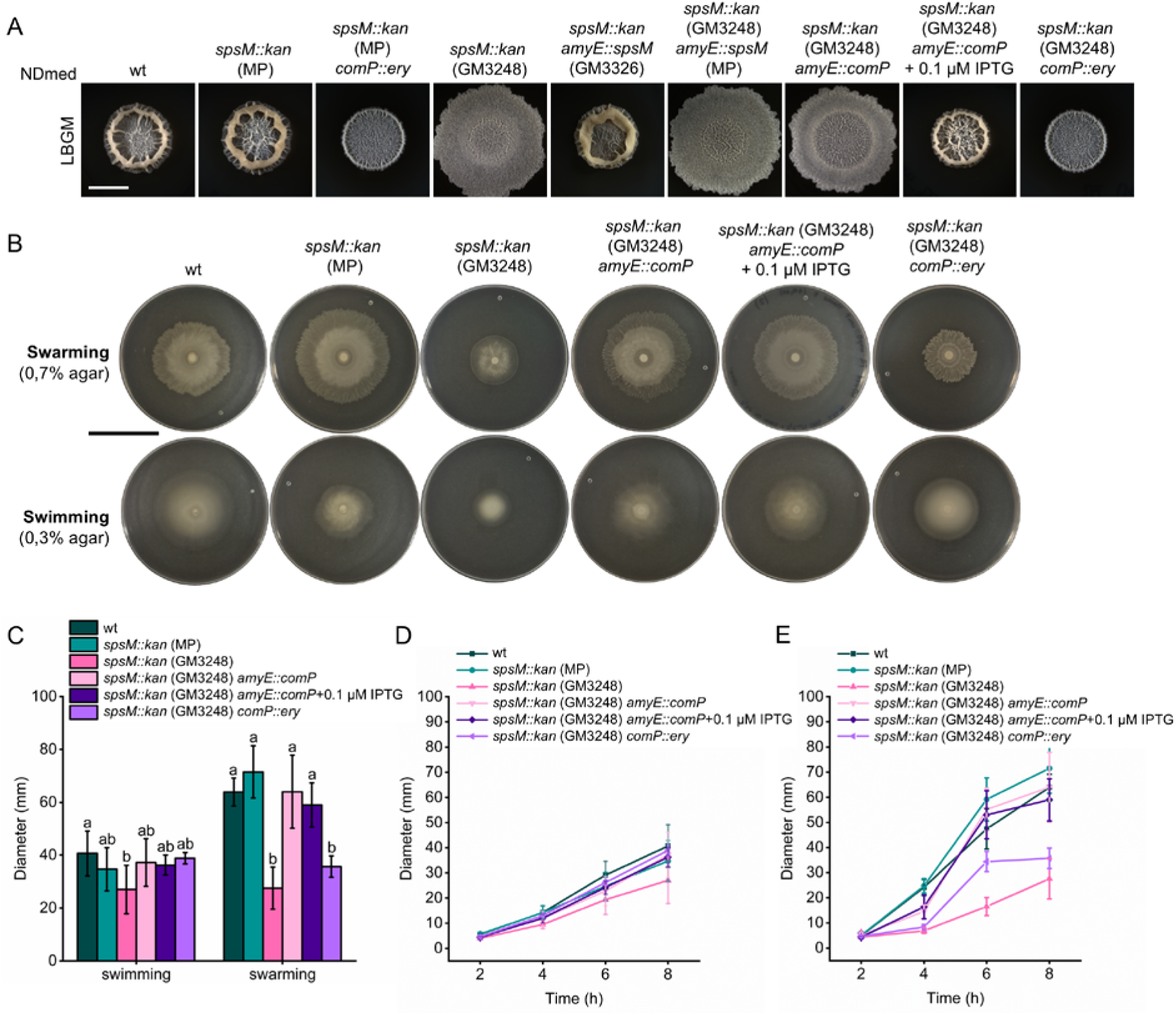
Impact of distinct *comP* mutations on biofilm morphology and motility. All strain labels refer to NDmed derivatives. The strains labeled as wt, *spsM::kan* (GM3248), and *spsM::kan amyE::spsM* (GM3326) were previously described by Sanchez-Vizuete *et al*. (2015). Strains marked with (MP) were constructed in this study. **A** Images of macrocolony biofilms of strains with and without an active *comP*. Macrocolony biofilms were grown on solid LBGM medium and imaged after 48 h incubation at 30°C. White scale bar: 5 mm. **B** Images of swarming and swimming motility of strains with and without an active *comP*. Plates were imaged after 8 h incubation at 37°C. Plates were imaged against a black background so that bacterial colonization zones appear white, and uncolonized agar appears dark. Swimming assays were performed on LB with 0.3% agar, and swarming assays on LB with 0.7% agar. Black scale bar: 5 cm. **C** Quantification of swimming and swarming diameters after 8 hours at 37°C. Columns represent mean diameters (n=6) and error bars represent standard deviation. Statistical analysis was performed using one-way ANOVA followed by Tukey’s Honest Significant Difference (HSD) test. Letters above error bars indicate statistically significant differences between strains separately for swimming and swarming (p < 0.05). **D, E** Dynamics of swarming (C) and swimming (D) motility over an 8-hour period, with measurements taken at 2 h, 4 h, 6 h, and 8 h.

The *comP* gene is part of the ComQXPA quorum sensing system in *B. subtilis* (Piazza et al., 1999; Weinrauch et al., 1990) and is involved in expression of genes involved in surfactin production (Msadek et al., 1991). Surfactin has been shown to enhance swarming motility in *B. subtilis* (Kearns & Losick, 2003; Kinsinger et al., 2003). To further confirm the impact of the spontaneous *comP* mutation in the NDmed *spsM::kan* (GM3248) strain, we compared motility phenotypes of wild-type and mutant strains on LB medium containing 0.3% agar (for swimming) and 0.7% agar (for swarming). A significant reduction in swimming motility was observed only in the NDmed *spsM::kan* (GM3248) strain compared to the wild-type but not compared to the rest of the tested strains (Figure 5B, 5C). The effect of the *comP* mutation was however more pronounced in swarming assays. Both the NDmed *spsM::kan* (GM3248) strain carrying the spontaneous *comP* mutation and the NDmed *spsM::kan* (GM3248) *comP::ery* deletion mutant exhibited reduced swarming motility compared to strains with an intact *comP* gene (Figure 5B, 5C). This reduction in motility was consistent across all measured time points, as shown in Figures 5D and 5E.

It has previously been shown that surfactin can be visualized as an expanding ring ahead of the swarming front, visible under reflected light (Julkowska et al., 2005). In line with this observation, strains lacking an active *comP* gene (NDmed *spsM::kan* (GM3248) and NDmed *spsM::kan* (GM3248) *comP::ery*) did not produce a visible surfactin ring during swarming. In contrast, the ring reappeared in the NDmed spsM::kan (GM3248) *amyE::comP* strain, where *comP* was complemented by an additional copy. Notably, this complementation restored both swimming and swarming motility, as well as surfactin production, even in the absence of IPTG, likely due to leaky promoter expression.

Interestingly, although the premature stop codon mutation (comP:c.1222T>A) and the complete deletion of *comP* produced similar swarming defects and lack of surfactin ring, they resulted in distinct macrocolony biofilm morphologies (Figure 5A), suggesting that these two mutations have different downstream effects on surface-associated phenotypes. Since bioinformatic analysis of the comP:c.1222T>A mutation predicts a secondary start codon located 15 bp downstream of the mutation it is possible both the N-terminal an C-terminal portion of ComP are separately translated and effect biofilm morphology in a distinct way.

### Permanent disruption of *spsM* promotes rapid emergence of spontaneous mutations affecting motility and biofilm morphology

In strains carrying the *spsM::kan* mutation two spontaneous mutations were identified: a deletion in *swrA* (c.26delT) in the P9_B1 fluffy strain and a substitution in *comP* (c.1222T>A) in the NDmed *spsM::kan* (GM3248) strain. These findings led us to suspect that disruption of *spsM* may accelerate the appearance of additional spontaneous mutations. To investigate this further, we first screened selected strains for spontaneous appearance of peripheral outgrowths. These outgrowths were restreaked and assessed for altered colony morphology (Supplementary Figure S5A).

Aside from flat colony morphotype similar to ‘fluffy’, we also observed an emergence of wider flat (similar to NDmed *spsM::kan* (GM3248) *comP* mutant) and a glossy colony types (Figure 6). A subset of representative strains from each morphotype was subjected to sequencing of *swrA* and *comP* loci. It turned out that all sequenced isolates with a shared morphology, carried the same mutation pattern within *swrA* and *comP*, allowing their classification based on phenotype. Strains labeled as mut-swrA displayed a flat and expanded colony morphology and carried the *swrA* c.26delT deletion. Mut-comP strain showed a wider, flatter morphology with reduced motility and carried a frameshift mutation in *comP*. Mut-unk strains exhibited a glossy colony appearance and altered swarming dynamics, though the exact mutation responsible was not identified (Figure 6, Supplementary Figure S5).

**Figure 6:**
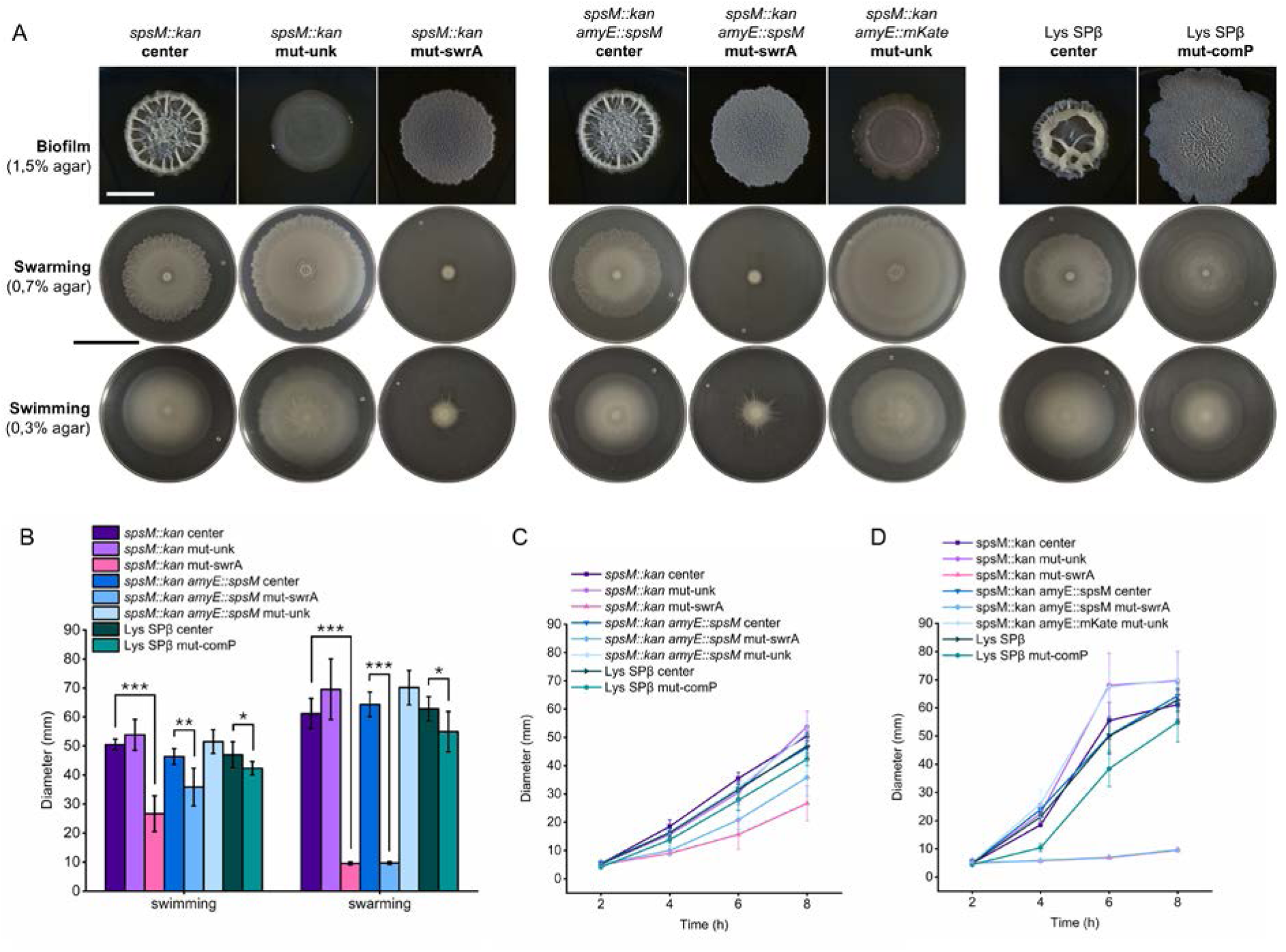
Impact of detected spontaneous mutations on biofilm morphology and motility. All strain labels refer to P9_B1 derivatives. **A.** Representative images of macrocolony biofilms, swarming motility, and swimming motility for spontaneous mutants. Strains were derived from the center (labeled as “center”) and outgrowth (labeled as “mut-”) regions of macrocolonies grown for 48 hours at 30 °C on solid LB medium. Mutants labeled as mut-swrA are associated with the *swrA* c.26delT deletion, mutant mut-comP carries the *comP* c.1115delT deletion, and mut-unk refers to strains with unknown mutations. Macrocolony biofilms were grown on LBGM agar and imaged after 48 hours at 30 °C. Swarming and swimming assays were performed on LB containing 0.7% and 0.3% agar, respectively, and imaged after 8 hours of incubation at 37 °C. Plates were imaged against a black background so that bacterial colonization zones appear white, and uncolonized agar appears dark. White scale bar: 5 mm. Black scale bar: 5 cm. **B** Quantification of swimming and swarming diameters after 8 hours at 37°C. Columns represent mean diameters (n=6) and error bars represent standard deviation. Statistical analysis was performed using one-way ANOVA followed by Tukey’s Honest Significant Difference (HSD) test, performed separately for each parental strain (*p < 0.05, **p < 0.01, ***p < 0.001). **C, D** Dynamics of swarming (C) and swimming (D) motility over an 8-hour period, with measurements taken at 2 h, 4 h, 6 h, and 8 h.

Despite potential variability between experiments, a consistent pattern emerged. No mutants were identified in the P9_B1 wt strain across all experiments. In contrast, spontaneous mutants described above, repeatedly appeared in *spsM::kan* and *spsM::kan amyE::spsM* strains. Interestingly, the one *comP* mutant (c.1115delT) emerged in the Lys SPβ background. Although its macrocolony morphotype resembled the NDmed *spsM::kan* (GM3248) strain carrying a different spontaneous *comP* mutation (c.1222T>A), the two strains displayed distinct swarm patterns. This suggests that different mutations in *comP* can lead to distinct effects on motility, even when overall colony morphology appears similar (Figure 6).

As additional controls, we tested 40 colonies of P9_B1 *amyE::mKate* and *spsM::kan amyE::mKate* strains. A single mutant appeared in the *spsM::kan amyE::mKate* strain, displaying the same glossy phenotype as the mut-unk group (Figure 6, Supplementary Figure S5). No mutations were observed in the *amyE::mKate* control. These results suggest that disruption of the *amyE* locus itself does not influence mutation emergence. Instead, the increased spontaneous diversification observed in *spsM::kan* and *spsM::kan amyE::spsM* strains indicate that the phenotypic/genotypic stability could not be restored by *spsM* complementation.

### Spontaneous point mutation in *swrA* provides a competitive advantage to strains with disrupted *spsM* in laboratory biofilm-promoting conditions

Among the spontaneous mutations identified in our screening, the most frequent was a single-base deletion in *swrA* (c.26delT). This mutation consistently occurred in P9_B1 *spsM::kan* strain at the same position and resulted in loss of swarming motility and a less complex biofilm morphology. Given that reduced biofilm complexity is commonly observed in laboratory strains of *B. subtilis* (Branda et al., 2001), we investigated whether the *swrA* c.26delT mutation, or a complete deletion of the gene (*swrA::tet*), might provide a fitness advantage in *spsM*-negative genetic background under laboratory biofilm-promoting conditions. To assess this, we performed competition assays on solid LBGM medium using 1:1 mixtures of fluorescently labeled strains. Cultures were incubated for 48 hours at 30 °C, and fluorescence was quantified by measuring integrated density from colony images (Figure 7). Because GFP and mKate fluorescent signals varied in intensity, monocultures were used to normalize the fluorescence values of cocultures (Figure 7A).

**Figure 7:**
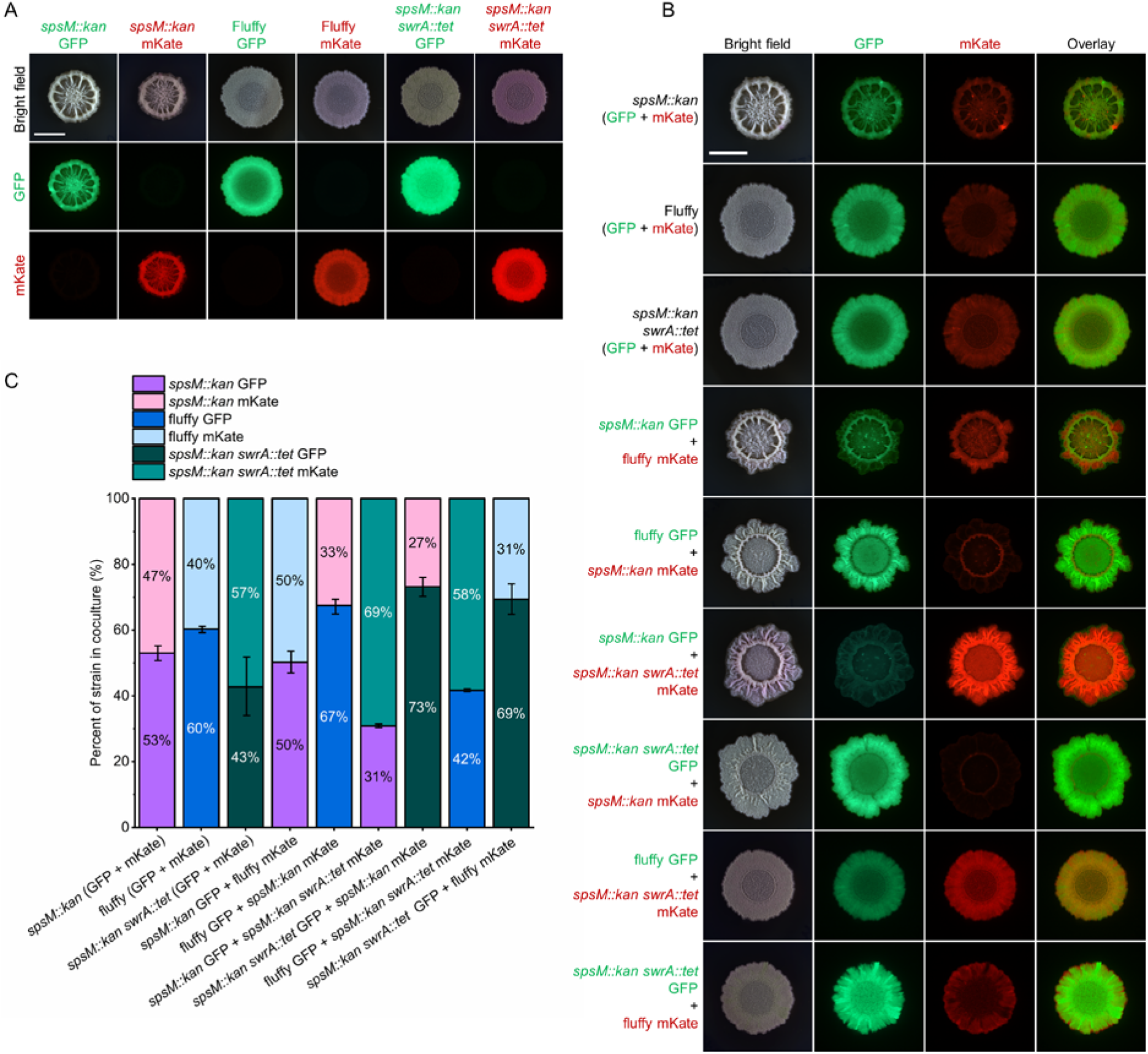
Competitive coculture assays comparing *swrA*-active and *swrA*-inactive strains in biofilm conditions. All strain labels refer to P9_B1 derivatives. The fluffy strain refers to the *spsM::kan* strain carrying the spontaneous swrA c.26delT mutation. **A, B** Representative macrocolony biofilms of monocultures (A) and 1:1 cocultures (B) grown on solid LBGM medium and incubated for 48 hours at 30 °C. Fluorescently labeled strains (GFP and mKate) were used to distinguish genotypes within each mixture. Scale bar: 5 mm. **C** Relative abundance of each strain in cocultures presented as a 100% stacked bar plot (n = 4). Fluorescence intensity was measured from macrocolony images using Fiji (ImageJ), and values were normalized using monoculture fluorescence to account for differences between GFP and mKate signal strength. Each bar shows the proportion of each strain in the coculture, reflecting its competitive fitness in biofilm-promoting conditions.

Fluorescent labels were first confirmed to have only a minimal impact on competitive fitness by comparing isogenic strains expressing either GFP or mKate. Next, the P9_B1 *spsM::kan* strain was mixed with the spontaneous mutant P9_B1 fluffy, which carries the *swrA* c.26delT mutation. In both labeling configurations, the fluffy strain showed a modest increase in relative abundance compared to the isogenic control, suggesting a slight competitive advantage (Figure 7B, 7C). When the *spsM::kan* strain was competed against the *spsM::kan swrA::tet* deletion strain, the effect was more pronounced. In both label combinations, the strain with the complete *swrA* deletion consistently outcompeted the strain with an intact *swrA* gene, indicating a clear fitness advantage under conditions with enhanced biofilm production. Interestingly, competition between the *spsM::kan swrA::tet* strain and the fluffy mutant again showed a competitive advantage for the deletion strain, even though both strains lacked a functional *swrA* gene. This mirrors the pattern observed for *comP*, where complete deletion had different phenotypic consequences compared to spontaneous point mutations.

In contrast, the same competitive advantage was not observed on LB medium alone (Supplementary Figure S6), indicating that the benefit of *swrA* deletion is condition dependent. These results suggest that loss of *swrA* function enhances fitness specifically under biofilm-promoting conditions on LBGM, likely by reducing the energetic cost of motility or regulatory interference. This may explain why the *swrA* c.26delT mutation was frequently recovered in our screen, as spontaneous mutations that promote growth under biofilm-favoring conditions are likely to be rapidly selected.

## DISCUSSION

The initial objective of our study was to gain deeper insight into the eco-evolutionary consequences of prophage integration at the functional *B. subtilis* locus *spsM*, previously linked to biofilm robustness (Sanchez-Vizuete et al., 2015). Our work took an unexpected turn as we observed a rapid accumulation of secondary loss-of-function mutations in *spsM::kan* strains. This suggests that *spsM* may either play a role in maintaining *B. subtilis* genomic stability, or that disruption of *spsM* imposes a strong selection for reduced biofilm formation and swarming motility.

The latter scenario would be analogous to the deletion of *ymdB*, which triggers the accumulation of suppressor mutations in *sinR*, where the initial deletion negatively impacts biofilm gene expression and the secondary mutations compensate for this effect by impairing the biofilm master repressor SinR (Kampf et al., 2018). A similar pattern was observed following deletion of the flagellin-encoding gene *hag*, which resulted in increased spontaneous diversification into a wrinkly colony morphotype, later linked to mutations in *sinR*. These secondary mutations compensated for the loss of motility by enhancing other multicellular behaviors that were more advantageous in the experimental setting (Richter et al., 2018).

In contrast to the above examples and previous reports (Kampf et al., 2018; Leiman et al., 2014; Richter et al., 2018), we did not observe any negative consequences of *spsM* loss that could potentially result in increased selective pressure. However, we did observe a clear fitness advantage of the secondary mutation in the *spsM*-negative background, which may instead indicate faster fixation of frequently occurring mutations (such as *swrA* loss-of-function mutation) in this background.

Loss-of-function mutations in key regulators are a common feature of microbial adaptation to laboratory conditions (a process referred to as domestication) and often target regulatory genes that govern energy-intensive traits, such as motility and biofilm formation. (Barreto et al., 2020; Dragoš, Lakshmanan, et al., 2018; Foster et al., 2015; Kampf et al., 2018; M. Martin et al., 2017; Richter et al., 2018; Schroeder et al., 2018; Sun et al., 2024). Similar patterns have also been reported in experimental evolution studies with *Pseudomonas*, *Escherichia*, and *Bacillus*, where adaptation to favorable laboratory conditions frequently results in loss-of-function mutations that downregulate redundant or costly processes (Dragoš, Lakshmanan, et al., 2018; Flynn et al., 2016; M. Martin et al., 2017; Rainey & Travisano, 1998; Traverse et al., 2013). Along these lines, in *B. subtilis*, mutations in *comP*, like mutations in *swrA*, affect multiple interconnected phenotypes and often confer a selective advantage in nutrient-rich media by eliminating costly communal behaviors (Barrick et al., 2009; Dragoš, Lakshmanan, et al., 2018; M. Martin et al., 2017) such as, collective motility, resulting in fitness advantage of mutations like *swrA* c.26delT.

The higher number of mutations we observed in *spsM::kan* mutants of *B. subtilis* is unlikely to be due to the loss of SpsM protein function per se, as complementation with a functional *spsM* gene at the *amyE* locus did not restore the phenotypic/genomic stability of the strain. This suggests that the increased genomic instability arises from structural consequences of the *spsM* locus disruption rather than the absence of protein. Disruption of *spsM* locus by a kanamycin resistance cassette may perturb local chromosomal architecture or regulatory elements, potentially interfering with chromatin organization, DNA replication, or recombination processes (Foster et al., 2015; Tenaillon et al., 2016). Additionally, the kanamycin cassette may exert a polar effect, where transcriptional read-through from the cassette activates downstream or neighboring genes. Notably, while *spsM* (∼2.1 Mb), *swrA* (∼3.5 Mb), and *comP* (∼3.1 Mb) are located on separate regions of the chromosome, trans effects via regulatory or physiological feedback loops cannot be excluded. Such local instability may propagate to more global mutation patterns, especially in the context of laboratory conditions that promote rapid growth and selection (Barrick et al., 2009; Brockhurst et al., 2011).

Interestingly, in case of SPβ lysogen, where *spsM* is interrupted by the prophage, only a single incident of spontaneous mutation was observed suggesting that, SPβ lysogens maintained a more stable genomic state compared to *spsM* deletion mutants This disparity may be explained by active lysogeny exhibited by the SPβ prophage, which can reversibly integrate and excise from the host genome, thereby buffering or obscuring the phenotypic consequences of *spsM* disruption (Abe et al., 2014; Rather et al., 2021). The SPβ integration site in *spsM* has been identified as a hotspot for dynamic prophage-host interactions, where reversible integration enables host cells to modulate phenotypes such as sporulation and biofilm formation in response to environmental cues (Abe et al., 2014; T. Dubois et al., 2020; Feiner et al., 2015; Sanchez-Vizuete et al., 2015). Interestingly, in certain temperate phages, integration events are very specific, targeted towards conserved, transcriptionally active sites (Feiner et al., 2015). It was proposed that such specific integration could represent an additional level of host control by the phage enabling prophages to hijack host regulatory circuits while minimizing deleterious effects (Abe et al., 2014, 2017; Butala & Dragoš, 2023; Feiner et al., 2015). While under experimental conditions the rapid emergence of *swrA* and *comP* mutations may be beneficial, the opposite may be true in a natural setting; therefore, prophage excision from *spsM* may benefit the host under natural conditions.

In our experiments, spontaneous mutations consistently emerged in *swrA* and *comP*, both of which are key regulators of motility and biofilm development (Kearns & Losick, 2003; Lin et al., 2025; Patrick & Kearns, 2012), underlining the significance of regulatory networks in bacterial evolution. The repeated emergence of *swrA* and *comP* mutations across independently evolved populations suggests strong selection pressure in nutrient-rich environments to suppress energetically expensive regulatory networks. Evolution in nutrient-rich environments tends to streamline genomes by selecting against energetically expensive, redundant, or non-essential traits (Barrick et al., 2009).

Mutations in *swrA* consistently led to a loss of swarming motility, aligning with its known function as a regulator of flagellar assembly regulation (Kearns & Losick, 2003; McLoon et al., 2011; Mukherjee & Kearns, 2014). Notably, reversible mutations in *swrA*, such as an A-T base pair insertion, have previously been described as arising via slipped-strand mispairing during replication (Calvio et al., 2005; Dubnau & Losick, 2006). The same frame-shift mutation is fixed in lab strains like *B. subtilis* 168, which lack swarming capability (Kearns et al., 2004; Patrick & Kearns, 2009). Beyond motility loss, we observed *swrA* mutations also altered biofilm morphology, likely due to downstream regulatory effects on exopolysaccharide and γ-poly-DL-glutamic acid (γ-PGA) synthesis, critical for robust biofilm formation (McLoon et al., 2011; Stanley & Lazazzera, 2005).

Similarly, mutations in *comP* impaired swarming motility, likely by abolishing surfactin production, which has been shown as a key surfactant required for surface spreading motility (Kearns & Losick, 2003). This is consistent with prior studies showing that ComP regulates surfactin and iturin biosynthesis through ComA phosphorylation, directly influencing both colony expansion and biofilm integrity (Kobayashi, 2007; Lin et al., 2025; Msadek et al., 1991; Sun et al., 2024). Our observations of surfactant-deficient colonies and altered morphology in *comP* mutants align with previously reported phenotypes in *B. subtilis* morphology (Julkowska et al., 2005; Kearns et al., 2004; McLoon et al., 2011; Sun et al., 2024). In addition to surfactin production, ComP-mediated signaling modulates other regulators like DegU, which further orchestrate motility and biofilm pathways (Amati et al., 2004; Msadek et al., 1991).

In light of previous reports on the role of spsM in biofilm formation (Sanchez-Vizuete et al., 2015) our results indicate that disruption of *spsM* alone (without secondary loss-of-function mutations) did not significantly alter host biofilm morphology or motility phenotypes. This observation was true for both the permanent disruption mutant (*spsM::kan*) and the SPβ lysogen, in which *spsM* is interrupted by prophage integration. These findings suggest a more refined understanding of *spsM* function and may call for a re-evaluation of the earlier hypothesis that *spsM* disruption promotes phage dissemination by restructuring biofilm spatial organization (Floccari & Dragoš, 2023; Sanchez-Vizuete et al., 2015).

On the other hand, our work revealed a novel insight into the functional and evolutionary implications of *spsM* disruption and therefore new interesting interplay between temperate phages and their host. Numerous studies have highlighted the profound impact of prophage carriage on host physiology, stress responses, and long-term evolutionary trajectories (Clokie et al., 2011; Howard-Varona et al., 2017; Mavrich & Hatfull, 2017; Stefanič et al., 2024). In *B. subtilis*, prophages have been shown to influence population dynamics and social behaviors, often conferring selective advantages such as enhanced competitiveness or altered developmental timing (Dragoš, Andersen, et al., 2021; Nanda et al., 2015). Our current findings extend this knowledge by highlighting a new role of prophage integration sites such as *spsM*, which may serve as evolutionary hotspots, where temperate phages exert long-lasting and multifaceted influences on host genome architecture and phenotypic diversity. The relative stability observed in SPβ lysogens emphasizes the adaptive advantage conferred by dynamic, regulated prophage integration, reinforcing the notion that prophage-host relationships are shaped by a balance of mutualism, parasitism, and co-evolution.

## ACKNWLOEDGMENTS

MP and AD were supported by Slovenian Agency for Research and Innovation (ARIS) grant no. P4-0116 and the European Union (ERC, PHAGECONTROL, 101041421). The views and opinions expressed are, however, those of the authors only and do not necessarily reflect those of the European Union or the European Research Council Executive Agency. Neither the European Union nor the granting authority can be held responsible for them. ID was supported by ARIS grants P4-0116 and J1-3021. TA was supported by an ARIS grant P4-0097. YD and RB were supported by supported by INRAE.

The stereomicroscopy experiments were supported by funding from ARIS to Infrastructural Centre Microscopy of Biological Samples (MRIC UL, I0-0022-0481-08), at Biotechnical Faculty, University of Ljubljana, Slovenia.

## SUPPLEMENTARY INFORMATION

**Figure S1.**
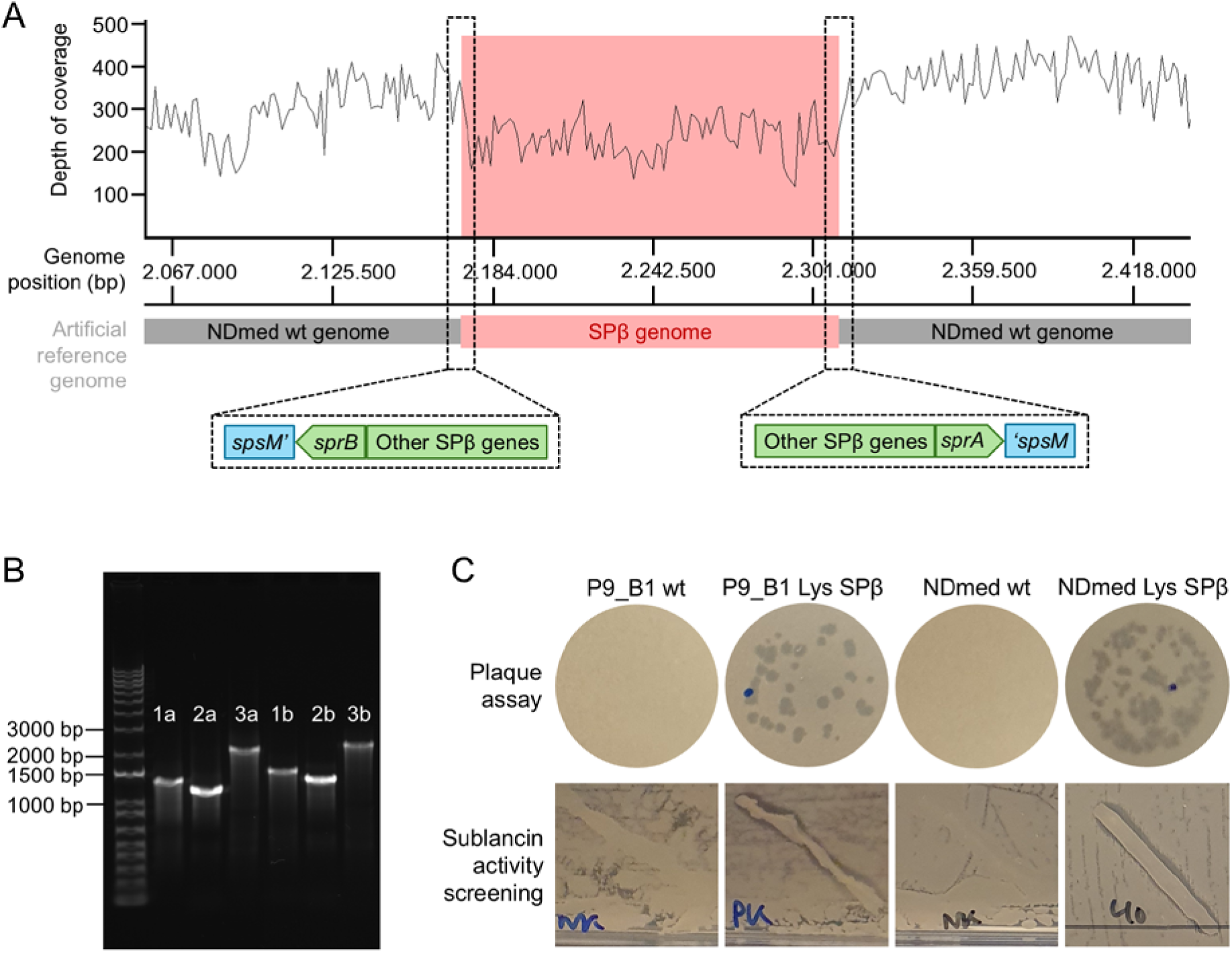
Confirmation of SPβ prophage integration into the *spsM* locus in NDmed and P9_B1 Lys SPβ strains. **A** Illumina sequencing reads from the NDmed Lys SPβ strain were aligned to an artificial reference genome in which the SPβ prophage sequence (NCBI Reference Sequence: NZ_CP045821.1) was inserted into the NDmed wild-type genome (GenBank accession: CP183238) at the expected *spsM* insertion site. The artificial genome was constructed in SnapGene. Read quality of Illumina reads was assessed using MultiQC, and reads were mapped to the artificial reference using Bowtie and SAMtools. Visualization of alignment coverage was performed in Artemis. The top panel shows the depth of coverage across the artificial genome. The lower schematic indicates genome structure, with the SPβ region highlighted in red and the flanking integration sites expanded to show the local gene organization. **B** PCR confirmation of SPβ integration in NDmed Lys SPβ and P9_B1 Lys SPβ strains. Letters “a” and “b” denote the NDmed and P9_B1 backgrounds, respectively. Lanes marked “1” show amplification of the left junction between *spsM* and SPβ, lanes “2” show the right junction, and lanes “3” indicate amplification of the sublancin gene cluster, a marker specific to the SPβ prophage. **C** Functional confirmation of SPβ prophage activity in lysogenized strains. Top row: Plaque assay showing SPβ phage release after mitomycin C induction. The appearance of plaques in the supernatant confirms the presence of inducible, active prophage. Bottom row: Sublancin-mediated antagonism assay. A 1:100 diluted lawn of the wild-type P9_B1 or NDmed strain was overlaid with a line of the lysogenized test strain. The presence of a clear inhibition zone around the line indicates sublancin production by SPβ-positive strains, confirming prophage functionality.

**Figure S2.**
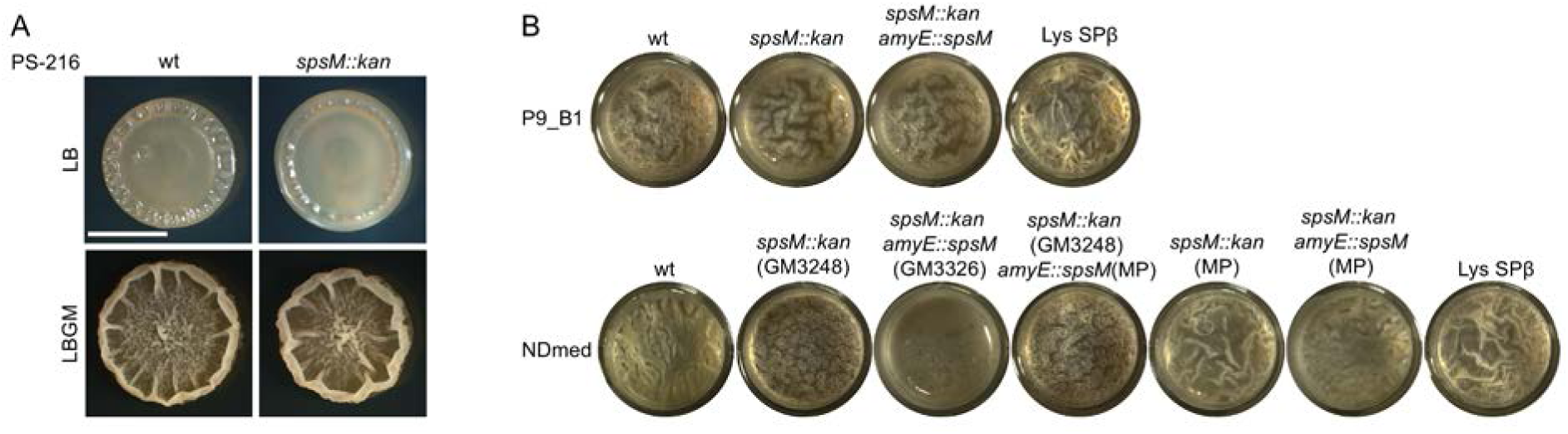
Macrocolony biofilm of PS-216 strains and pellicle morphology of *B. subtilis* P9_B1 and NDmed strains. **A** Macrocolony biofilm morphology of PS-216 strains grown on LB and LBGM media at 30°C for 48 hours. Scale bar: 5mm **B** Pellicle biofilm morphology of P9_B1 and NDmed strains grown statically in liquid LBGM medium at 30°C for 48 hours.

**Figure S3.**
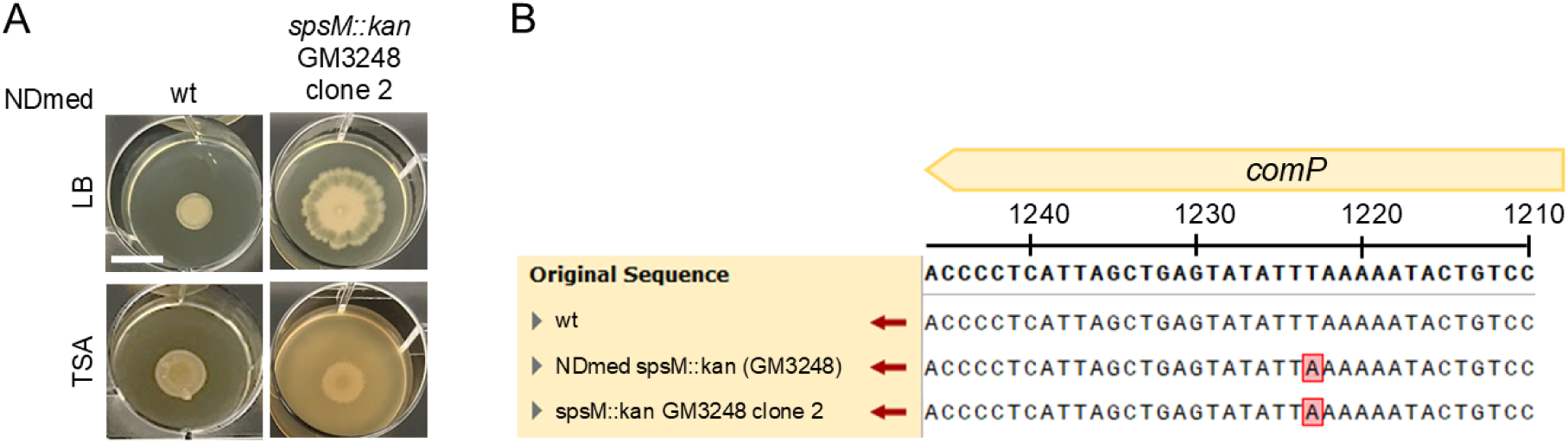
Macrocolony morphology and *comP* gene sequences of original NDmed *spsM::kan* mutant stock. **A** Macrocolony biofilm morphology of original NDmed *spsM::kan* GM3248 clone 2 stock grown on LB and TSA media at 30 °C for 5 days. Scale bar: 10 mm. **B** Sanger sequencing of the targeted region in the *comP* gene. The top panel shows a schematic representation of the wild-type *comP* reference sequence (“Original Sequence”). Below, Sanger sequencing results from each strain are aligned and compared to the reference. Deviations from the wild-type sequence are highlighted with a red box. NDmed spsM::kan (GM3248) label refers to the bacterial stock stored at Biotechincal faculty, University of Ljubljana, and spsM::kan GM3248 clone 2 label refers to the original bacterial stock made at the Micalis Institute (Sanchez-Vizuete et al., 2015).

**Figure S4.**
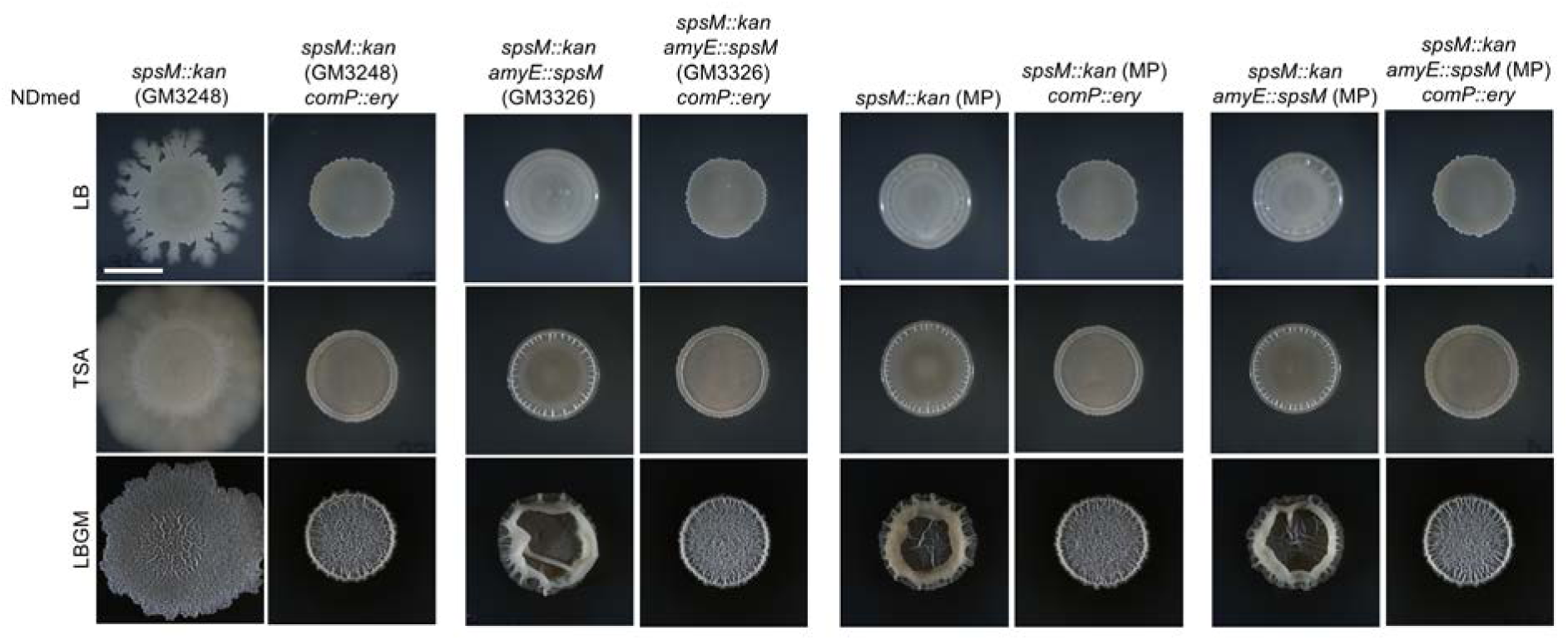
Macrocolony morphology of NDmed strains with and without *comP::ery* deletion. Macrocolonies were grown on LB, TSA, and LBGM media at 30°C for 48 hours. Scale bar: 5 mm

**Figure S5.**
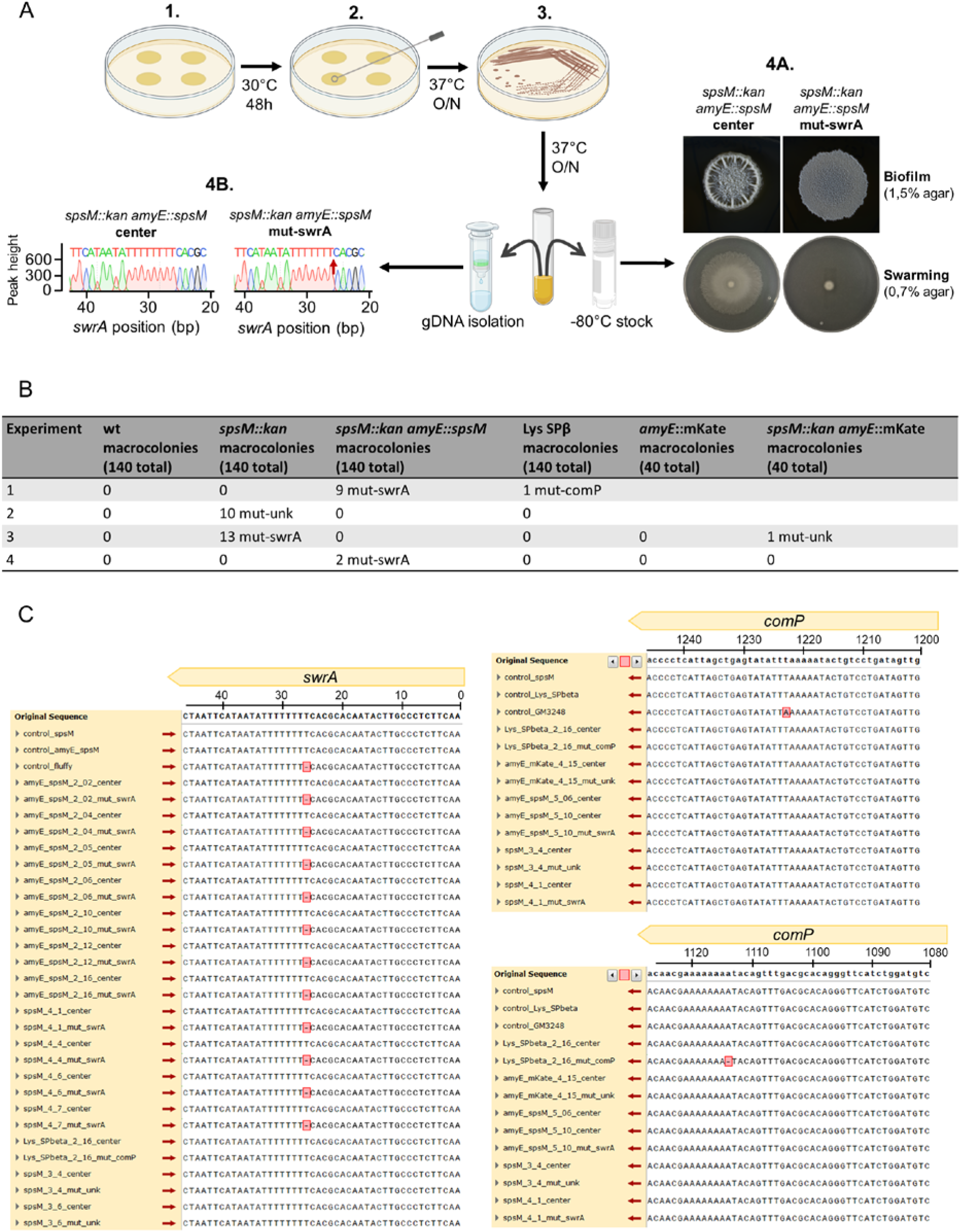
Experimental workflow and results of spontaneous mutation screening in strains with active or disrupted *spsM*. A Schematic overview of the experimental setup. All strains were inoculated on the same LB agar plate, with each plate representing one biological replicate (1). Macrocolonies were incubated for 48 hours at 30 °C. If peripheral outgrowths appeared, both the center and outgrowth of the same macrocolony were transferred to fresh LB plates (2). After further incubation, colony morphology was compared (3). Outgrowths with altered morphology were considered potential spontaneous mutants. A single colony from both the center and the outgrowth was selected to prepare –80 °C glycerol stocks and isolate genomic DNA (gDNA). Stocks were later used to assess biofilm morphology and motility phenotypes (4A), and the gDNA was used to amplify the *swrA* and *comP* genes for Sanger sequencing (4B). **B** Summary of number of outgrowths with altered morphology across four independent experiments, classified by originating strain and phenotype classification. Only a subset of mutants were sequenced (C). Nevertheless, all sequenced isolates sharing a morphological phenotype were confirmed to harbor identical mutations, validating the use of phenotype-based classification. Strains labeled mut-swrA showed colony morphology identical to those confirmed to carry the *swrA* c.26delT deletion. The single mut-comP strain carried the *comP* c.1115delT deletion. The mut-unk phenotype was characterized by a glossy colony morphology and altered swarming dynamics; however, the specific underlying mutation remains unidentified. **C** Sanger sequencing data of targeted regions in the swrA and comP genes. At the top of each aligment, schematic representations indicate the wild-type reference sequences (labeled as “Original Sequence”). Below are aligned nucleotide sequences from each analyzed strain. Strains are labeled according to their origin: colonies derived from the central region of macolonies are labeled “center,” and colonies from peripheral outgrowths are labeled “mut-”. Background genotypes are indicated as spsM (*spsM::kan*), amyE_spsM (*spsM::kan amyE::spsM*), and Lys_SPbeta (Lys SPβ). Strains labeled “control” represent the original parental stocks utilized for mutation screening. Additionally, sequences from the fluffy and GM3248 strains are presented as controls, representing the initial mutations identified, which motivated further mutation screening efforts. Sequence deviations relative to the wild-type are highlighted by red boxes.

**Figure S6.**
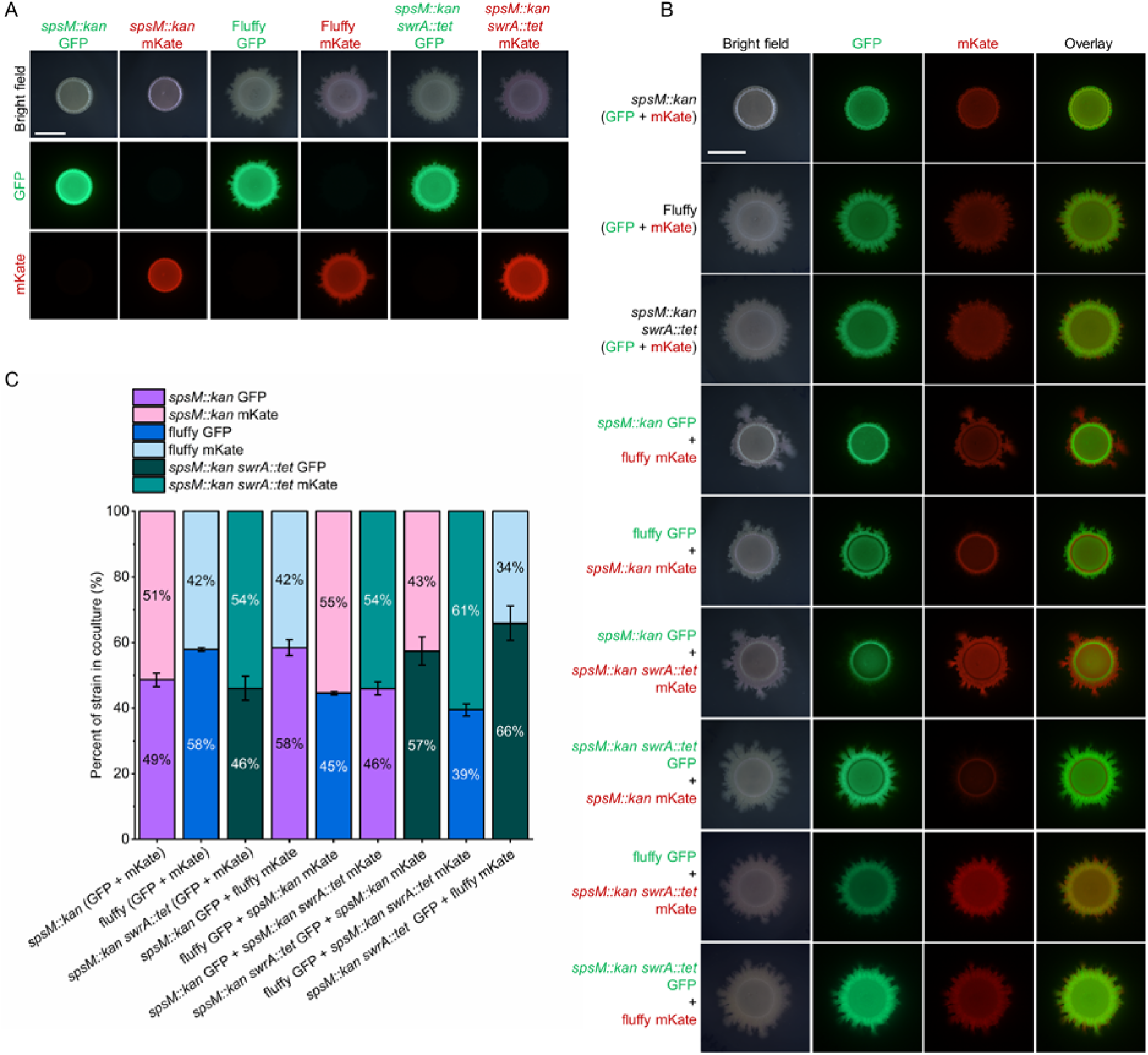
Competitive coculture assays comparing *swrA*-active and *swrA*-inactive strains grown on LB medium. All strain labels refer to P9_B1 derivatives. The fluffy strain refers to the *spsM::kan* strain carrying the spontaneous *swrA* c.26delT mutation. **A, B** Representative macrocolonies of monocultures (A) and 1:1 cocultures (B) grown on solid LB medium and incubated for 48 hours at 30 °C. Fluorescently labeled strains (GFP and mKate) were used to distinguish genotypes within each mixture. Scale bar: 5 mm. **C** Relative abundance of each strain in cocultures presented as a 100% stacked bar plot (n = 4). Fluorescence intensity was measured from macrocolony images using Fiji (ImageJ), and values were normalized using monoculture fluorescence to account for differences between GFP and mKate signal strength. Each bar shows the proportion of each strain in the coculture, reflecting its competitive fitness in biofilm-promoting conditions.

